# LicD-Mediated Cell Wall Decoration Governs Phage Sensitivity in Enterococcus faecalis Clinical Isolates

**DOI:** 10.1101/2025.05.13.653012

**Authors:** Muhammed Awad, George Bouras, Sholeh Feizi, Susanna R. Grigson, Peter-John Wormald, Alkis J. Psaltis, Sarah Vreugde

## Abstract

*Enterococcus faecalis* has emerged as a prevalent antibiotic-resistant pathogen in clinical settings. Herein, we report the identification of three novel lytic phages targeting vancomycin-resistant *E. faecalis*. While the isolated phages all belonged to the *Kochikohdavirus* genus, there were distinctive differences in their tail fibre proteins, affecting their adsorption. The phages showed strong antibacterial activity with wide host range, infecting >90% of the tested *E. faecalis* clinical and hospital wastewater isolates (n=13) with variable efficiency. The variation in host range was genomically correlated to the presence of the *LicD* gene in phage sensitive bacteria, which is responsible for phosphorylcholine decoration of the bacterial cell wall. Furthermore, the isolated phages were predicted to harbour genes encoding for depolymerase enzymes, which was confirmed by *in vitro* testing showing a >80% reduction in biofilm biomass. Phages inhibited bacterial growth for ≥12 hours, followed by the emergence of bacteriophage insensitive mutants (BIMs) that were 4-fold more sensitive to vancomycin compared to parent strains. In addition, the isolated BIMs showed less capability of evading THP-1 macrophage and produced weaker biofilms. These findings underpin the potential of the isolated phages in combating recalcitrant *E. faecalis* associated biofilm-mediated infections.

## Introduction

*Enterococcus faecalis* is a leading example of the world’s antimicrobial resistance crisis. *E. faecalis* was previously considered as a commensal Gram-positive anaerobe with negligible clinical significance [1]. It is now known to be a prevalent pathogen in nosocomial infections [2]. *E. faecalis* is involved in the pathogenesis of various medical conditions including failure of endodontic treatment [3], urinary tract infections [4], wound infections [5] and endocarditis [2]. *E. faecalis* has multiple strategies to combat antibiotics including intrinsic resistance to several antibiotics such as penicillins, cephalosporins, aminoglycosides and vancomycin [6]. In addition, they can acquire antimicrobial resistance genes from neighbouring cells through pheromone sensitive plasmids [7]. Furthermore, *E. faecalis* can shift their growth mode to become sessile aggregated colonies covered with extracellular polymeric substances known as biofilms [2]. When grown in biofilms, bacteria can become up to 1000 times more tolerant to antibiotics, which leads to infection recurrence and treatment failure [8]. *E. faecalis* biofilms have specifically been linked to endodontic diseases and implant failure in the oral cavity [9, 10]; infesting medical devices and causing nosocomial infections. The versatility of antibiotic resistance approaches implemented by *E. faecalis* require the use of unconventional treatment options to augment the fight against this pathogen.

Phage therapy is among the unconventional approaches that has been revived recently to overcome antibiotic-resistance [11]. Since bacteriophages depend solely on bacterial cells for their survival, they are continuously evolving bacterial targeting techniques [12]. After encountering target bacteria, a phage reversibly adsorbs to a bacterial surface receptor via specific proteins in the phage’s tail i.e. receptor binding proteins [13]. After the primary adsorption, the phage is irreversibly attached to the bacterial surface through a secondary receptor and starts injecting its DNA into bacterial cell to replicate, followed by assembly of phage particles and release of new phages from bacteria using phage encoded enzymes [14]. The phage adsorption can be assisted by phage lytic enzymes such as depolymerases to solubilize the polysaccharide capsule shielding the bacterial surface [15]. Phages that can produce depolymerase enzymes are considered especially valuable assets, as they can disrupt the biofilm matrix, overcoming phenotypic resistance of biofilms [15, 16].

The literature contains many studies showcasing the potential of lytic phages in inhibiting the growth of antibiotic-resistant E. *faecalis* [17, 18]. However, like antibiotics, bacteria can quickly become resistant to a single phage, limiting wide clinical application [14, 19]. Thus, isolating and combining new phages that can target different bacterial receptors into versatile phage cocktails is considered a promising strategy to limit the occurrence of phage resistance during the course of treatment [19]. To date, *E. faecalis* phage infection protein and exopolysaccharides have been identified as targets for bacteriophages [18]. Therefore, isolating new phages and studying their potential receptors would benefit the formulation of more versatile phage cocktails. In this study, we report the isolation and characterization of novel bacteriophages targeting multi-drug-resistant *E. faecalis* isolated from different sources. We showcase the genetic versatility of *E. faecalis* genomics and report potential new receptors targeted by *E. faecalis* phages.

## Materials and Methods

### Bacterial strains collection and identification

The host bacterium used for phage isolation is *Enterococcus faecalis* ATCC 700802 (American Type Culture Collection; ATCC, Manassas, VA, USA). Clinical *E. faecalis* isolates from patients with diabetic foot infection (n = 4) and chronic rhinosinusitis (CRS) (n =2) were collected using bacterial swabs after obtaining informed consent with protocols approved by the Central Adelaide Local Health Network Human Research Ethics Committee in Adelaide, South Australia reference number (HREC/15/TQEH/132). Swabs were placed in a sterile Amies transport medium (Sigma Transwab, MWE Medical Wire, Corsham, UK) and transported to the laboratory for processing. Further *E. faecalis* strains (n = 7) were isolated from wastewater samples obtained from the Queen Elizabeth Hospital, Adelaide, SA. Briefly, 100 µL of the collected samples were spread on Difco™ m Enterococcus selective Agar (Thermo Fisher Scientific Australia Pty Ltd., Scoresby, VIC, Australia) and incubated at 37°C. After 24 hours, red colonies were isolated and streaked on Brain Heart Infusion (BHI) agar (Oxoid, Thebarton, SA, Australia); uniform colonies were selected and identified using MALDI-TOF mass spectrometry (Bruker, Billerica, USA).

### Antibiotic sensitivity testing

The sensitivity of the isolated bacteria to vancomycin and ampicillin (Sigma Aldrich, St. Louis, MO, USA) were determined through the minimum inhibitory concentration (MIC) using the broth microdilution assay [20]. Briefly, overnight cultures of bacterial strains were adjusted to 0.5 McFarland followed by 1:100 dilution in cation adjusted Muller Hinton broth. Two-fold dilutions of vancomycin and ampicillin were prepared in the same media and mixed with bacterial suspensions. MIC was determined as the minimum concentration that inhibited bacterial growth.

### Bacterial whole genome sequencing

Bacterial DNA was extracted using DNeasy blood & tissue kit (Qiagen, Hilden, Germany) according to the manufacturer’s protocols. The extracted DNA was hybrid sequenced for all isolates other than 2 clinical isolates (DFI250 and DFI266) and isolated BIMs, where only long-reads were available. Long-read sequencing was conducted in house using the Oxford Nanopore Technologies (ONT) platform on a MinION Mk1B (Oxford Nanopore Technologies, Oxford, UK) device. Specifically, the Rapid Barcoding Kit (Oxford Nanopore Technology, SQK-RBK 114.96) was used to sequence *E. faecalis* whole genomes on R10.4.1 MinION flow cells (Oxford Nanopore Technology), using 50 ng of the isolated DNA. Base-calling was conducted with Dorado v0.5.0, using the r10.4.1_e8.2_400bps_sup@v4.2.0 basecaller model (Oxford Nanopore Technology).

Short-read sequencing was conducted on the Illumina platform, using the Illumina NextSeq 550 (Illumina Inc, San Diego, USA) and NextSeq 500/550 Mid-Output kit v2.5 (Illumina Inc., FC-131-1024) at SA Pathology (Adelaide, SA, Australia). To prepare for short-read sequencing, the genomic DNA was isolated using the DNeasy blood & tissue kit (Qiagen, Hilden, Germany) according to manufacturer’s instructions. The sequencing libraries were prepared using a modified protocol for the Nextera XT DNA library preparation kit (Illumina Inc. FC-131-1024). The genomic DNA was fragmented, after which a low-cycle PCR reaction was used to amplify the Nextera XT indices to the DNA fragments. Sequencing yielded one hundred fifty bp paired end reads.

### Bacterial genome assembly and annotation

Bacterial genomes were assembled with a long-read first approach supplemented with short-read polishing using Hybracter v0.7.3 [21]. Specifically, long-reads were filtered with Filtlong v0.2.1 [22]. Long-read adapters were trimmed using Porechop_ABI v 0.5.0 [23], while short-reads were quality controlled using fastp v0.23.4 [24]. Long-reads were assembled using Flye v2.9.3 [25]. The bacterial chromosomes (i.e. circular contigs above 2.5MB) were extracted and reoriented to begin with the dnaA chromosomal replication initiator gene with Dnaapler v0.8.0 [26]. Long-read polishing was skipped with the ‘--no_medaka’ parameter, as it has recently been shown to add no improvements for Dorado SUP model reads [21]. Short-read polishing was then conducted using Polypolish v 0.6.0 [27] followed by Pypolca v0.3.1[28]. Plassembler v1.6.2 [29] was used to assemble all plasmids using both read sets.

All complete bacterial genomes were annotated using Bakta v1.9.4 with the full database [30]. Antimicrobial gene detection was also conducted with AMRfinderplus v4.0.3 [31]. A pangenome was generated using Panaroo v1.5.1 [32], while plasmids were typed using the mob_typer command from Mob Suite v3.1.9 [33].

### Bacteria Insensitive Mutants (BIMs) Bioinformatics Analysis

A consensus high quality ATCC 700802 genome sequence was generated with Trycycler v0.5.4 [34]. To evaluate the potential of phage sequences to map to homologous sections of the bacterial genome, the ATCC 700802 genome sequence was combined with the three phage genomes to create a master reference genome sequence. Each BIM read set was mapped against this master reference. Reads were mapped with Minimap2 v2.27. Variants were then called using Clair3 v1.0.10 [35] as the best performing Nanopore variant caller [36]. Variants with an alternative allele frequency of 70% or higher were kept as first pass. Following this, variants were screened and visualised manually to determine if they were likely to be real, or artifacts caused by systematic errors in Nanopore sequencing (such as in long homopolymer regions). A full set of visualised pileup plots can be found in the Supplementary Figures.

### Phage Isolation and purification

Wastewater samples obtained from Glenelg and Bolivar wastewater treatment plants in Adelaide, SA, Australia, were centrifuged at 10,000×g at 4 °C for 30 minutes; the supernatant was collected, and filter sterilized using a 0.22 µm syringe filter (PALLAcrodisc. NY, USA). Phages were isolated using a double agar layer method as previously described [37]. Briefly, 500 µL of wastewater were mixed with 100 µL of *E. faecalis* ATCC 700208 overnight culture for 10 minutes followed by mixing with 4 mL of 0.4% W/V BHI molten agar; poured to 1.5% W/V BHI agar plates and incubated at 37°C. After 24 hours, clear plaques were picked using sterile pipette tips and stored in 500 µL SM buffer (100 mM NaCl, 8 mM MgSO4, and 50 mM Tris-HCl (pH 7.5)) for further purification. Phage purification was performed using double layer agar (DLA) method using collected phage lysates. Plaques with similar morphology were collected in SM buffer. The process was repeated three times to ensure the purity of the isolated phages. A single plaque from the third round was picked and used for further amplification and characterization.

### Phage amplification, purification and storage

A single colony of *E. faecalis* ATCC 700802 was grown in 50 mL BHI broth. At mid-log phase OD = 0.2 equivalent to 3 × 10^8^ CFU/mL, phage lysates were added and incubated at 37 °C for 3-6 hours. Chloroform was added at 1:100 ratio and allowed to incubate at 37 °C for 5 minutes.

The culture was centrifuged at 4000 ×g for 10 minutes to remove bacterial debris; the lysate without chloroform was filter sterilised using 0.2 µm syringe filters and plaque forming units (PFU) were determined using DLA method. To extract the phages from crude lysates, precipitating solution (10% PEG-8000, 1 M NaCl final concentration) was mixed with phage lysates in 1:2 ratio precipitant: lysate with gentle swirling and kept at 4 °C overnight. The precipitated lysate was centrifuged at 10,000 ×g for 30 minutes. The supernatant was discarded, and phage pellets were resuspended in SM buffer. To remove PEG, the lysate solution was mixed with chloroform in a 1:1 ratio; gently mixed and centrifuged at 10,000 ×g for 10 minutes at 4 °C. The aqueous layer containing the pure phage was collected and stored at 4 °C for further testing.

### Phage Genome sequencing assembly and annotation

Phage DNA was extracted using (Norgen Biotek Crop, Thorold, ON, Canada) extraction kit according to the manufacturer’s instructions. The phage genome was sequenced using long read sequencing platform (MinION Nanopore sequencer) after preparing sequencing libraries using Oxford Nanopore rapid barcoding kit 96 V14 (SQK-RBK114.96) on R10.4.1 MinION flow cells (Oxford Nanopore Technology), using 50 ng of the isolated DNA. according to the manufacturer protocol with a run time of 48 hours.

Phage genomes were assembled and annotated using Sphae v1.4.2 [38]. Specifically, long-reads were filtered with Filtlong v0.2.1 [22]. Long-read adapters were trimmed using Porechop_ABI v0.5.0 [23]. Long-reads were assembled using Flye v2.9.3 [25]. The phage genomes were then annotated using Pharokka v1.7.3 [39] using Phanotate v1.5.1[40] as the gene caller. Annotations were then improved using Phold v0.2.0 [41]. Specifically, protein structures were generated for each predicted protein using Colabfold v1.5.5 [42] using the commands ‘colabfold_search’ (using the UniRef30 and Colabfold environmental databases for multiple sequence alignment generation) and ‘colabfold_batch --num-models 5’. The top-ranking model was selected as the structure for each protein. Following this, Foldseek v9.427df8a [43] was used to compare phage protein structures to a database of Colabfold [42] generated phage protein structures based on the PHROGs database [44]. Taxonomic assignment was performed by comparing the phage genomes to their closest Mash hit [45] to the INPHARED database [46].

DepoScope [47] was used to predict depolymerase genes in each phage. Specifically, both binary classification and per-token output predictions were considered. LoVis4u v0.0.7 [48] was run on the Phold output to generate genome synteny plots. The homo-oligomeric state of the depolymerase genes was determined by re-running Colabfold with AlphaFold-multimer [49] with varying chain copy numbers. Specifically, the multiple sequence alignments generated using colabfold_search were modified to generate structures for N=2-9 chains. These modified alignments were then used to generate structures with the command ‘colabfold_batch –num-models 3 –num-recycle 5 --model-type alphafold2_multimer_v3’. For each protein, the structure with the highest mean interface predicted template modelling (ipTM) score, mean predicted local distance difference test (pLDDT) and minimum pDockQ2 score [50]was used to infer the most probable oligomeric state [51].

### Bacteriophage adsorption rate

Bacteriophage adsorption rate was determined as previously described with slight modifications [52]. Briefly, overnight culture of *E. faecalis* ATCC strain was adjusted to 1 × 10^8^ CFU/mL followed by mixing with different phages at titre of 1× 10^6^ PFU/mL in BHI media and allowed to stand for 5 minutes at 37 ⁰C without disturbance. Control tubes containing BHI media with no bacteria was incubated with equal titres of phages to determine the total phage concentrations (P0). Following 5 minutes, the tubes were immediately transferred to ice bath followed by centrifugation at 6000 ×g for 5 minutes at 4 ⁰C to precipitate bacteria with the adsorbed phages. A 100 µL of supernatant from both tubes were collected and free phage titre (P) was enumerated using double layer agar assay (DLA); the concentration of the bacteria (B) in 1 mL was determined via CFU counting. The adsorption rate constant (K) was quantified using the following equation

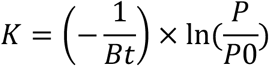

The experiment was performed on three different days and the average of the adsorption rate constant was reported.

### One step growth curve

Phage growth kinetics were determined using one step growth curve as previously described [53]. Each bacteriophage was mixed with *E. faecalis* at MOI 0.01 in BHI broth and allowed to adsorb on bacteria for 5 minutes followed by centrifugation at 6000 ×g for 5 minutes to pellet the infected bacteria; supernatant with unabsorbed phages was discarded and the pellets were diluted 1:100 in BHI broth and incubated at 37 °C under mechanical shaking at 180 rpm. Aliquots of 100 µL were taken at different time points and serially diluted to determine phage titres using DLA method. The experiment was repeated thrice, and the average phage titres were plotted against different time points to determine the burst size and latent period. Burst size was calculated by dividing the average of maximum phage titre after plateau/ initial phage titre, and the latent period is determined as the period after which phage titre increased.

### Determination of host range

The host range of the isolated phages was tested as previously described [54, 55] using 13 *E. faecalis* clinical isolates in addition to the host bacterium. Moreover, the specificity of these bacteriophages to *E. faecalis* was evaluated against a collection of Gram-positive bacteria (n =11) including *E. faecium*, *S. aureus*, *S. lugdunensis* and *S. epidermidis* **(Supplementary Table S3).** Briefly,100 µL of bacterial overnight culture were mixed with 0.4% molten BHI agar and poured on 1.5% BHI agar plates and air dried for 10 minutes at room temperature; 5 µL of each phage lysates at 10^6^ PFU/mL were spotted on bacterial lawns in triplicates and 5 µL of SM buffer was spotted in the middle of the plate as a control.

### Determining the efficiency of plating (EOP)

Efficiency of plating (EOP) was determined as previously described [55, 56]. Bacteriophage stocks were serially diluted followed by inoculating 20 µL of phage lysates with 100 µL of overnight bacterial cultures and 0.4% molten BHI agar and poured into BHI agar plates. Following 16-18 hours incubation at 37 °C, the plaque forming units (PFU) were enumerated and EOP was calculated as follows:

EOP = average PFU on tested bacteria /average PFU on host bacteria

EOP (0.5-1.0) is considered as high efficiency; EOP 0.2-0.5 moderate efficiency; 0.001-0.2 low efficiency; inefficient < 0.001 [55].

### Determination of thermal and pH stability

The stability of phages at various pH and temperatures were determined as previously described [54]. For thermal stability, 1 mL of phage stocks (10^9^ PFU/mL) was incubated at various temperatures between 4 °C and 100 °C for one hour followed by determination of phage titre. For pH stability, aliquots of phage stocks were diluted 1:10 in SM buffer at a range of pH values between (2-12) and incubated for 1 h at room temperature followed by quantification of the phage titres using DLA method.

### Transmission Electron Microscopy (TEM)

The phages morphology was characterized using FEI Tecnai G2 Spirit 120KV Transmission Electron Microscopy. 5 µL of purified phage stock (10^9^ PFU/mL) were placed on the carbon coated side of carbon/formvar coated grid (ProSciTech Pty Ltd., Townsville, QLD, Australia). The sample was allowed to attach on the coated grid for 3 min followed by mixing with 5 µL of fixative solution (1.25% glutaraldehyde, 4% paraformaldehyde in phosphate buffer solution (PBS) and 4% sucrose) for 2 minutes. Finally, 5 µL of 2% uranyl acetate were added and allowed to stand for 2 min before capturing images using TEM. For capturing the adsorption of phages to *E. faecalis* cell wall, bacterial suspension was incubated with bacteriophage at MOI 10 and incubated for 30 minutes followed by centrifugation for 5 minutes at 1000xg force. The pellets were collected and suspended in fixative solution and TEM images were captured.

### Bacteriophage time-kill assay and isolation of bacterial insensitive mutants (BIMs)

To investigate the antibacterial kinetics of the isolated Phages against *E. faecalis*, time-kill assay was conducted using different multiplicity of infections (MOIs). Briefly, overnight culture of *E. faecalis* ATCC 700802 was adjusted to OD_600_ 0.2 equivalent to 1.5× 10^8^ CFU/mL followed by 1:100 dilution in BHI broth. Different bacteriophages individually and in 1:1 cocktail were mixed with bacterial suspension at different MOI in the range between (100 - 0.01) with a final volume of 200 µL. The plate was incubated in CLARIOstar^®^ plate reader at 37 °C and absorbance measurements at 600 nm were recorded hourly for 24 hours. The readings were corrected to negative controls (media only). Aliquots of turbid wells were collected and spread on BHI agar plates and the sensitivity of the obtained colonies against different phages and Phage cocktails were assessed using DLA method. Isolates that were confirmed resistant to phage infection were termed BIMs.

### Assessment of BIMs fitness to evade phagocytosis

BIMs uptake by macrophages and inactivation were evaluated using THP-1 cell line and compared to the parent ATCC strain. Briefly, THP-1 cells (ATCC, Manassas, USA) were seeded at 0.15 × 10^6^ cells in 750 μL RPMI containing 200 ng/mL phorbol 12-myristate 13-acetate (PMA) (Sigma-Aldrich, St Louis, USA) in a 24-well plate and incubated at 37 °C for 48 h in a fully humidified incubator at 5% CO_2_. Cells were infected with *E. faecalis* ATCC-700802, BIMs 10, 16 and 20 at MOI 50 and 10 for 1 and 4 h for uptake and inactivation studies respectively. Cells exposed to media only were considered a negative control and bacteria incubated in media were used as bacterial infection control. Supernatants were collected and the remaining extracellular bacteria were removed using lysostaphin (10 µg/mL) (Sigma-Aldrich) for 15 min. Cells were lysed with 0.1% triton X-100 for 30 min to assess intracellular infection. Serial dilution of supernatants and cell lysates was prepared in 0.9% saline and plated on BHI agar plates. Colony forming units (CFUs) were enumerated after overnight incubation at 37 °C. Experiments were repeated 3 times.

### Assessment of BIMs ability to form biofilms

The ability of the generated BIMs to form biofilms were assessed using crystal violet assay. Briefly, aliquots of overnight culture of *E. faecalis* ATCC 700208 and BIM 10, 16 and 20 were adjusted to 1 McFarland equivalent to 3 × 10^8^ CFU/mL followed by 1:15 dilution in BHI broth. Aliquots of 180 µL of bacterial suspension were placed in 96 well plate (BMG Labtech, Ortenberg, Germany) and allowed to grow statically for 24 h at 37 °C. The grown biofilms were washed with 0.9% saline and stained with 0.1% W/V crystal violet stain. The absorbance of the stained biofilms was recorded at 595 nm and the average of three independent assays was recorded.

### Determination of antibiofilm activity

The antibiofilm activity of the isolated phages against established biofilms was determined using crystal violet assay. *E. faecalis* biofilms were established on 96 well plates as previously described [57]. Briefly, aliquots of overnight culture of *E. faecalis* ATCC 700208 was adjusted to 1 McFarland equivalent to 3 × 10^8^ CFU/mL followed by 1:15 dilution in BHI broth. Aliquots of 180 µL of bacterial suspension were placed in 96 well plate (BMG Labtech,

Ortenberg, Germany) and allowed to grow statically for 24 h at 37 °C. The obtained biofilms were washed twice with saline solution to remove unattached cells before adding the treatment. 180 µL of phage lysates in the range between 10^9^-10^5^ PFU/mL were added to the biofilms and untreated wells containing media only served as controls. Following 24 hours of incubation, the biofilms were washed and fixed using methanol; air dried and stained with 0.1% crystal violet (Sigma-Aldrich, Castle Hill, NSW, Australia) for 20 minutes. The plates were rinsed with distilled water and air dried. Crystal violet was solubilized using 33% v/v acetic acid (Chem-Supply, Adelaide, SA, Australia) and the absorbance was measured at 595 nm using FLUOstar Optima microplate reader (BMG Labtech, Ortenberg, Germany). The percent biofilm biomass reduction was calculated in comparison to the control samples.

## Statistical analysis

Statistical calculations were performed using Graph Pad Prism v 9.0.0 (121) (GraphPad Software, San Diego CA, USA). The Independent Samples t-test was used to assess the statistical differences between the means of two groups. The results were considered statistically significant when the p-value was <0.05. *p < 0.05; **p < 0.01; ***p < 0.001 and ****p < 0.0001.

## Results

### Isolation of three novel phages targeting vancomycin resistant *E. faecalis*

The aim of this study is to isolate and characterize bacteriophages with potential clinical application against vancomycin resistant *E. faecalis.* Therefore, we used the ATCC strain 700802, which is vancomycin resistant as a host for phage isolation. Thereafter, we determined the susceptibility of a panel of *E. faecalis* strains to both ampicillin and vancomycin, commonly used antibiotics to control Enterococcal infections [58, 59]. Although the tested strains were isolated from different sources (clinical and hospital wastewater samples), they all showed high resistance to both antibiotics with MIC values above the breakpoints [60] except for Efa-W6 that was sensitive to vancomycin and the ATCC strain that was sensitive to ampicillin (**Supplementary Table S1**).

Three *Enterococcus* phages were isolated from two different sites of wastewater treatment plants in Adelaide at different time points. The phages were named APTC-Efa.10, APTC-Efa.16 and APTC-Efa.20.

### Bacteriophages showed strong activity against clinical isolates and moderate to no activity against wastewater isolated bacteria

Initially, the spotting technique was conducted to determine the sensitivity of the isolates to the phages. Clear plaques were considered sensitive, while turbid plaques were considered as semi sensitive strains [54, 56] (**Table S2**). All phages showed clear zones on the clinical isolates except CI-1104 that was only semi-sensitive to APTC-Efa.16 and APTC-Efa.20. In contrast, the ability of the phages to infect *E. faecalis* isolated from the hospital wastewater was limited; APTC-Efa.10 could not infect most of the wastewater isolated *E. faecalis* except for Efa-W2 and Efa-W8 while APTC-Efa.16 and 20 showed turbid plaques on all wastewater isolates. We extended the testing of the host range against other Gram-positive cocci including *E. faecium*, *S. aureus*, *S. epidermidis* which all showed resistance to the isolated phages (**Table S3).**

To confirm the competency of phages against *E. faecalis*, efficiency of plating (EOP) was calculated to determine the ability of phages to produce high progeny, which is an important aspect for phage therapy [55, 56]. The EOP data was in line with the spot test assay, where high productive infection (EOP ≥ 0.5) was observed against all clinical isolates for all three phages except for clinical isolate CI-1104 where EOP between 0.1-0.3 was seen. In contrast, the phages showed moderate to weak productivity against most wastewater samples (EOP <0.5) (**Table1).**

### Morphology, phage kinetics and stability studies show all 3 phages belong to the *Myoviridae family,* efficiently propagate and are stable at various temperatures and pH

TEM images depicted that all 3 phages have long contractile tails characteristic of *Myoviridae* family with tail length ∼ 200 nm and icosahedral capsid with a diameter of ∼89 nm (**Figure 1)**.

**Figure 1.**
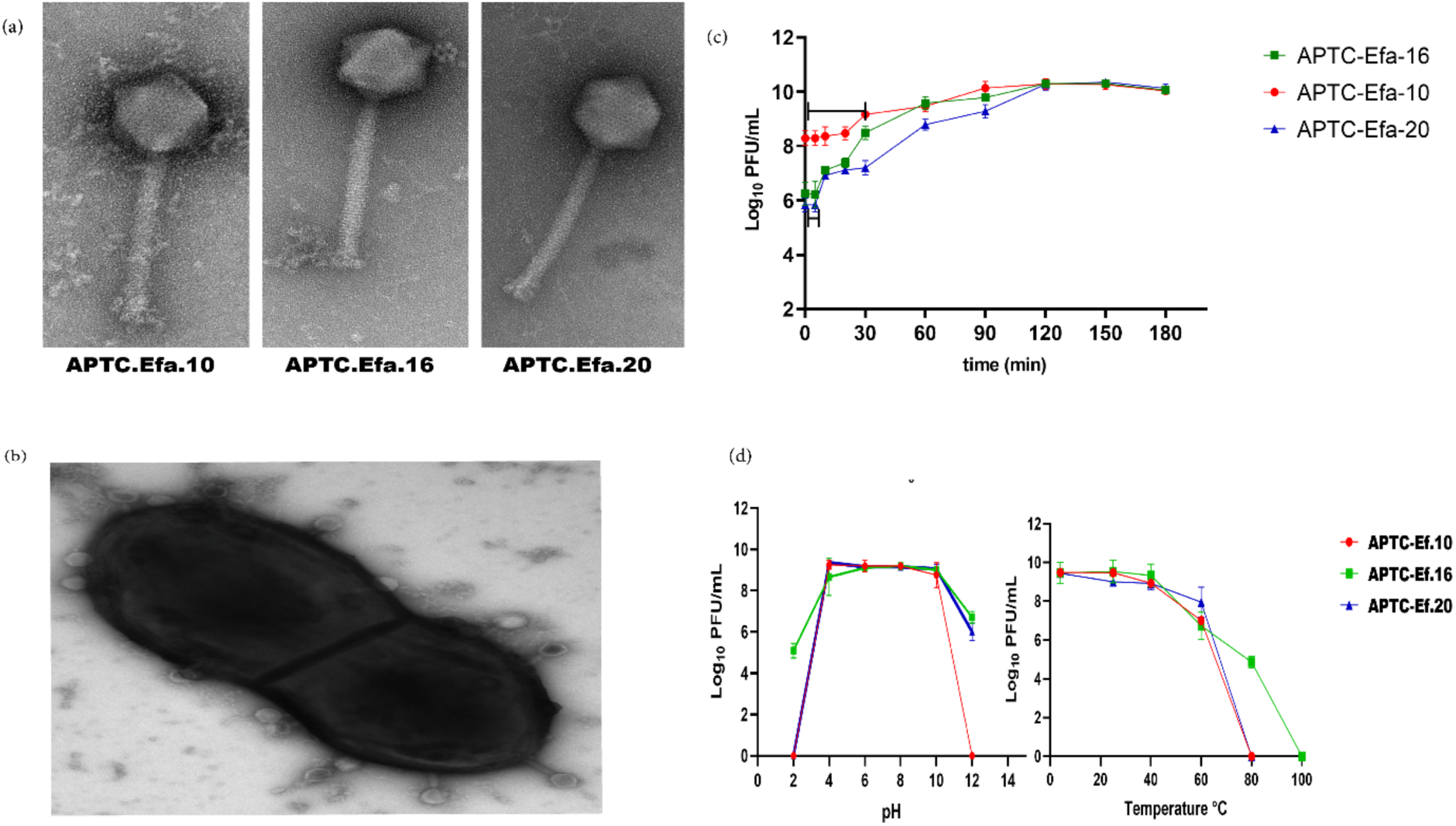
*Characterization* of *E. faecalis* phages. (a) TEM images depicting icosahedral heads and contractile tails identical to *Myoviridae* family; (b) TEM image showing APTC-Efa.16 infecting *E. faecalis* cell; (c) one step growth curve of the 3 phages; (d) pH and temperature stability of the isolated phages.

Among the three phages, APCT-Efa.16 showed the fastest adsorption rate k= 1.1 × 10^-10^ mL/min followed by APTC-Efa.20 k= 9 × 10^-11^ mL/min and APTC-Efa.10 K= 4 × 10^-11^ mL/min. The one step growth curve of the isolated phages showed some similarities between APTC-Efa.16 and APTC-Efa.20 with a latent period of 10 minutes. APTC-Efa.10 depicted a longer latent period of 30 minutes. The rise period of the 3 phages peaked at 120 minutes followed by a plateau indicating the end of the first infection cycle. APTC-Efa.16 scored the highest burst size with 45 PFU/cell followed by APTC-Efa.20 and APTC-Efa.10 with burst sizes of 29 and 22 PFU/cell respectively.

The resilience of phages to changes in pH and temperature was then determined. There was no significant change in the viability of the isolated phages between pH 4 and 10. However, APTC-Efa.16 was the most tolerant to pH changes recording 4 log_10_ reductions in viability at pH 2 compared to the physiological pH value 7 and ∼3 log_10_ reductions in viability at alkaline pH 12. Likewise, the viability of APTC-Efa.20 was reduced by 3 log_10_ at pH 12 but this phage could not survive acidic pH 2 (**Figure 1 d**). On the other hand, APTC-Efa.10 could not survive pH values of 2 and 12, however viability was unchanged between pH 4 and 10. In terms of thermal stability, the viability of all phages did not change between 4 and 40 °C, while the decline in viability started at 60 °C leading to complete inactivation of APTC-Efa.10 and APTC-Efa.20 at 80 °C. In contrast, APTC-Efa.16 was the most tolerant to high temperatures with 50% of the phages surviving incubation at 80 °C and inactivation occurring only at 100°C (**Figure 1 d**). These data depict that the isolated phages are stable at regular pH and storage temperatures, which are desirable characteristics for their usage in a therapeutic context [61].

### Structure-based phage annotation suggests variation in baseplate wedge and depolymerase-containing tail fibres

Next, we looked at the phage genomes. Taxonomic classification suggested that all three phages are closely related, with all belonging to the *Herelleviridae* family and the *Kochikohdavirus* genus. All three phages had guanine-cytosine content percentage (GC%) of around 36 % and coding sequence density of approximately 91%. Their lengths ranged from 138.5 kbp (APTC-Ef.20) up to 145 kbp (APTC-Ef.16) (**Supplementary Table S3**). Genome annotation with Pharokka and Phold indicated the absence of antimicrobial resistance (AMR) and integration-excision genes in all three phages, suggesting a strictly lytic-lifestyle (**Table S4**). Interestingly, all 3 phages possess a structural homolog to the *cll* gene from the gop-beta-cll anti-phage defense system that is likely to be a part of a toxin-antitoxin system [62].

Broadly, gene synteny was preserved between the three phages (**Figure 2 a).** Interestingly, there were 2 structural (i.e. proteins annotated as head and packaging, connector or tail) proteins with the same predicted function that showed significant variation. The first of these was annotated as a baseplate protein by Pharokka and Phold (coloured blue in **Figure 2 a)**. The lengths varied from 1131 residues for phage APTC-Ef.10, 804 residues for phage APTC-Ef.16 and 467 residues for phage APTC-Ef.20. Extra structural based annotation using predicted Colabfold structures, Foldseek webserver analysis and manual searches against PDB 4v96 [63] suggest that this protein is a baseplate upper protein that connects the tail fibres to the phage tail. Oligomeric state prediction of this protein indicated that in APTC-Efa.10 and 16, it is most likely to form monomers, while in APTC-Efa.20 forms a trimer that each connect the distal tail end to a tail-fibre protein. (**Supplementary Figure S1**).

**Figure 2.**
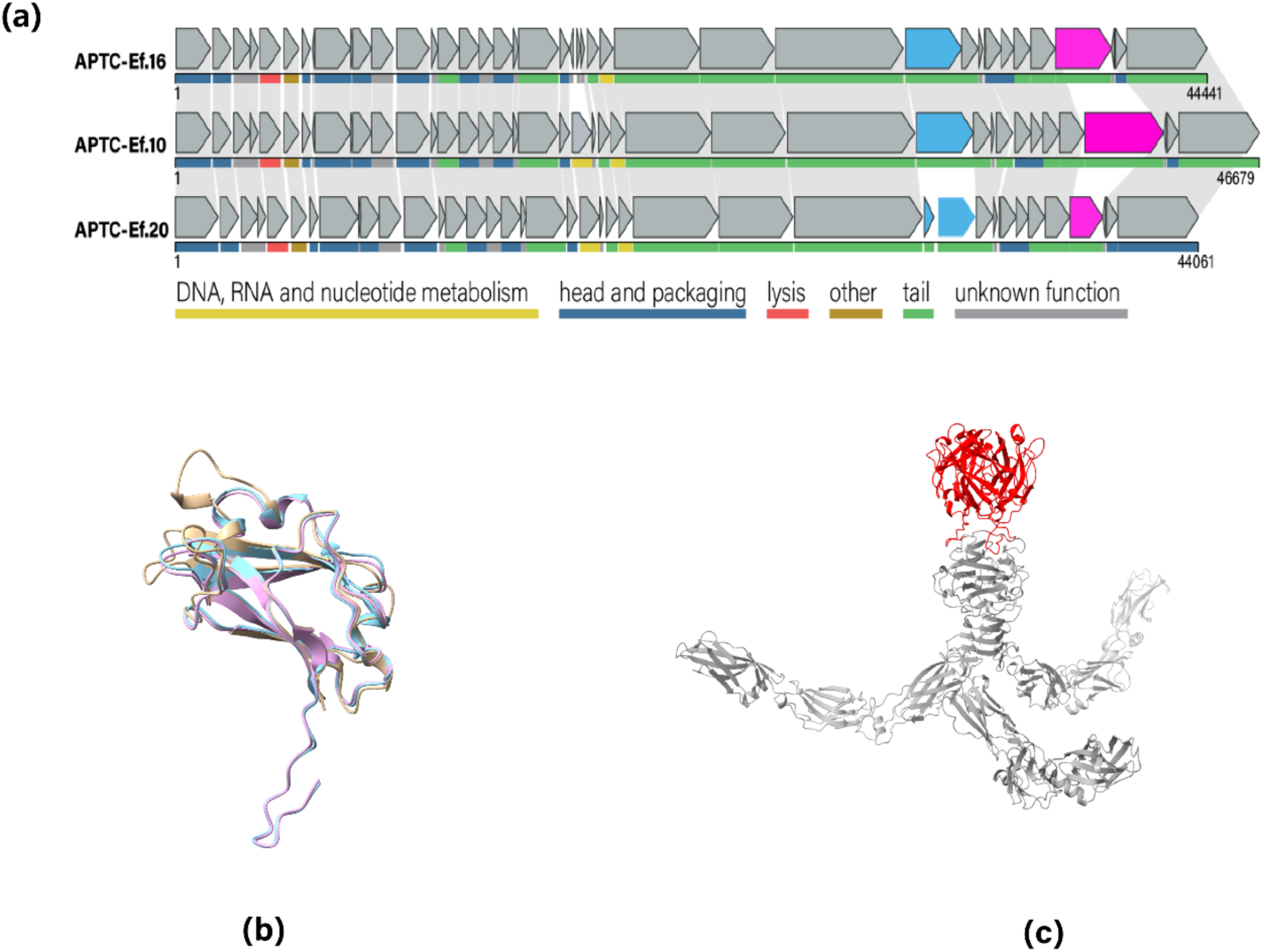
*(a) Gene synteny plot for the structural genes in APTC-Ef.10, APTC-Ef.16 and APTCEf.20*. All genes were reoriented to begin with the predicted large terminase gene. Similar genes are linked by grey shaded regions, while unrelated genes have no shading. The bright pink colour represents the variable baseplate proteins, while blue colour represents the predicted tail fibre proteins with predicted depolymerase domain. Each gene function is coloured beneath by its predicted PHROG category function; *(b) Three depolymerase domains in APTC-Ef.10 (gold), APTC-Ef.16 (blue) and APTC-Ef.20 (pink) as predicted by Colabfold; (c) Protein structure of the tail fibre trimer from APTC-Ef.20.* Red represents depolymerase domains that combined to form a “drill-like” spike, while the grey colour represents the remainder of the tail fibre.

The other structural protein that significantly varied between the three phages was annotated as a “tail fibre protein” (coloured pink in **Figure 2 a)**. DepoScope analysis suggested that these proteins likely carried a beta-propeller depolymerase domain. Specially, APTC-Efa.16 was extremely likely (>99% likelihood) to possess a beta-propeller depolymerase domain from residues 685 to 805 in CDS 30. APTC-Ef.20 also shared a very similar domain in CDS29, while APTC-Efa.10 did not have a depolymerase domain that was detected by DepoScope (**Figure S2**). However, combining the protein structural prediction and comparison methods using Colabfold and Foldseek revealed that APTC-Efa.10 in fact did possess a structurally near-identical depolymerase domain (**Figure 2 b and Figure S2**). Oligomeric state analysis also revealed these tail fibres containing depolymerase domains formed trimers, with the depolymerase domains from each copy combining to form a “drill-like” spike (**Figure 2 c**). The depolymerase activity of these phages was then further elaborated experimentally by testing their ability to disrupt *E. faecalis* biofilm matrix.

### Phages have potent antibiofilm activity against *E. faecalis* biofilm

The antibiofilm activity of individual phages against established biofilms was then probed. Firstly, the minimum concentration of phages that can eradicate biofilms (MBEC) (defined as a reduction in biomass of > 80% compared to control) was determined using the ATTC-700802 host. Biofilm biomass reduction of ≥ 80% was achieved after treatment with titres of 1×10^9^ and 1×10^8^ PFU/mL similar to previous reports with Enterococcus phages [64, 65](**Figure 3a**). Thereafter, we used MBEC of 1×10^8^ PFU/mL to test the antibiofilm activity of phages against two *E. faecalis* clinical isolates from patients with diabetic foot infections (CI 8010 and CI 8050). These isolates were selected because all 3 phages showed consistent activity against both isolates with EOP ≥0.9. All phages significantly reduced the biofilm biomass of both isolates (p<0.001) with more than 80% reduction in OD_595_ readings (**Figure 3b**).

**Figure 3.**
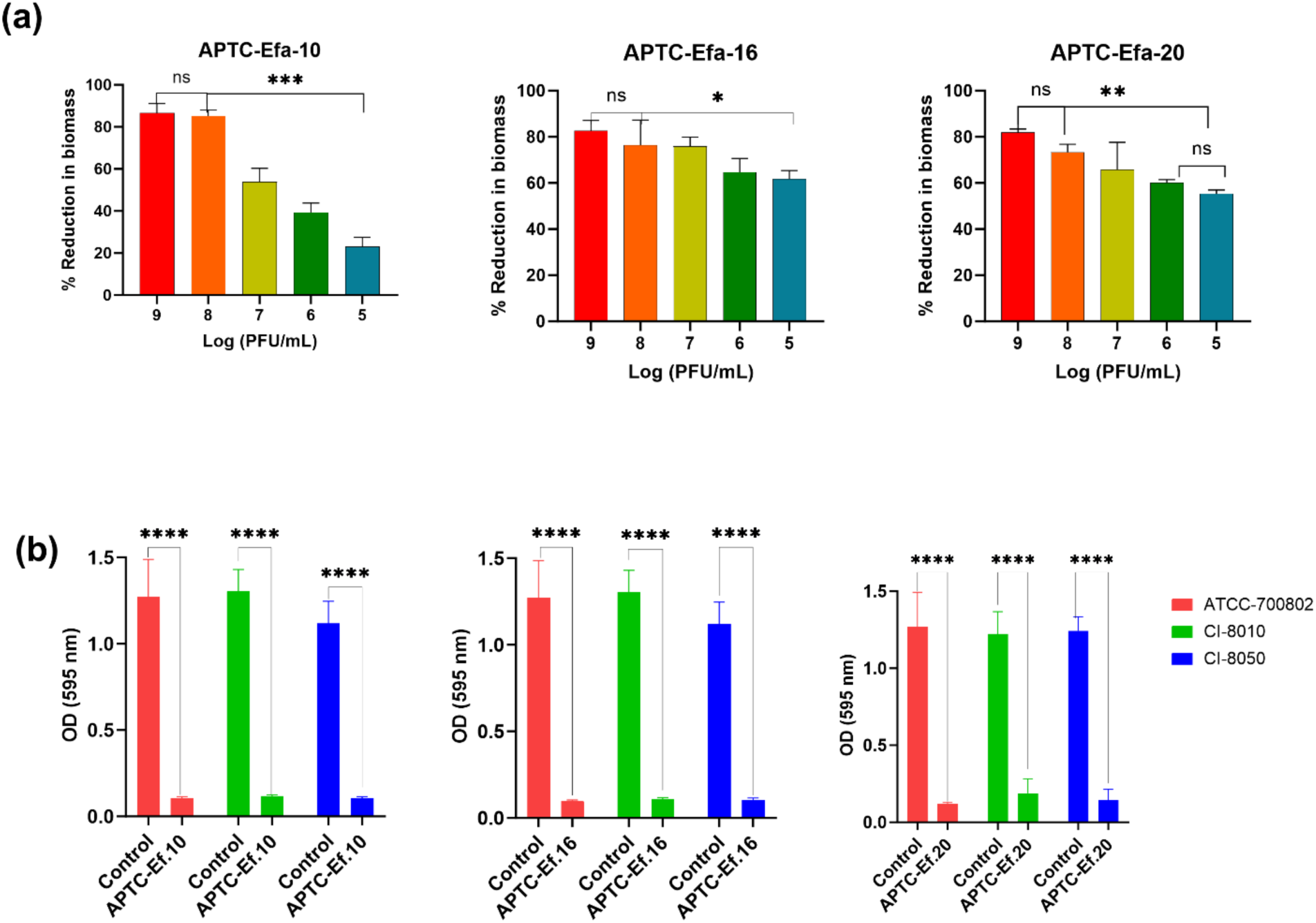
*Antibiofilm activity of Enterococcus phages*. (a) Minimum biofilm eradication concentration presented as percent reduction in the biofilm biomass compared to non-treated control using phage titres ranging from (9-5) log_10_ PFU/mL. (b) antibiofilm activity of Enterococcus phages against host ATCC-700802 and clinical isolates 8010 and 8050. Data presented as mean ± SD n=3. ns: non-significant, p > 0.05, *p < 0.05; **p < 0.01, ***p < 0.001 one-way ANOVA followed by multiple comparisons tests.

Given the predicted presence of depolymerase encoding genes in the phage’s tail fibres, we tested the potential of the two phages with strongest antibiofilm activity (APTC-Ef-16 and APTC-Ef-20) against *E. Faecalis* isolated from wastewater where both phages showed poor to moderate infections as interpreted from EOP data ranging between (0.3 and 0.01) (Table 1). Interestingly, both phages could reduce 50-60% of biofilms biomasses despite the poor antibacterial activity against planktonic cells probably due the depolymerase activity of those phages (**Figure S3**).

**Table 1.**
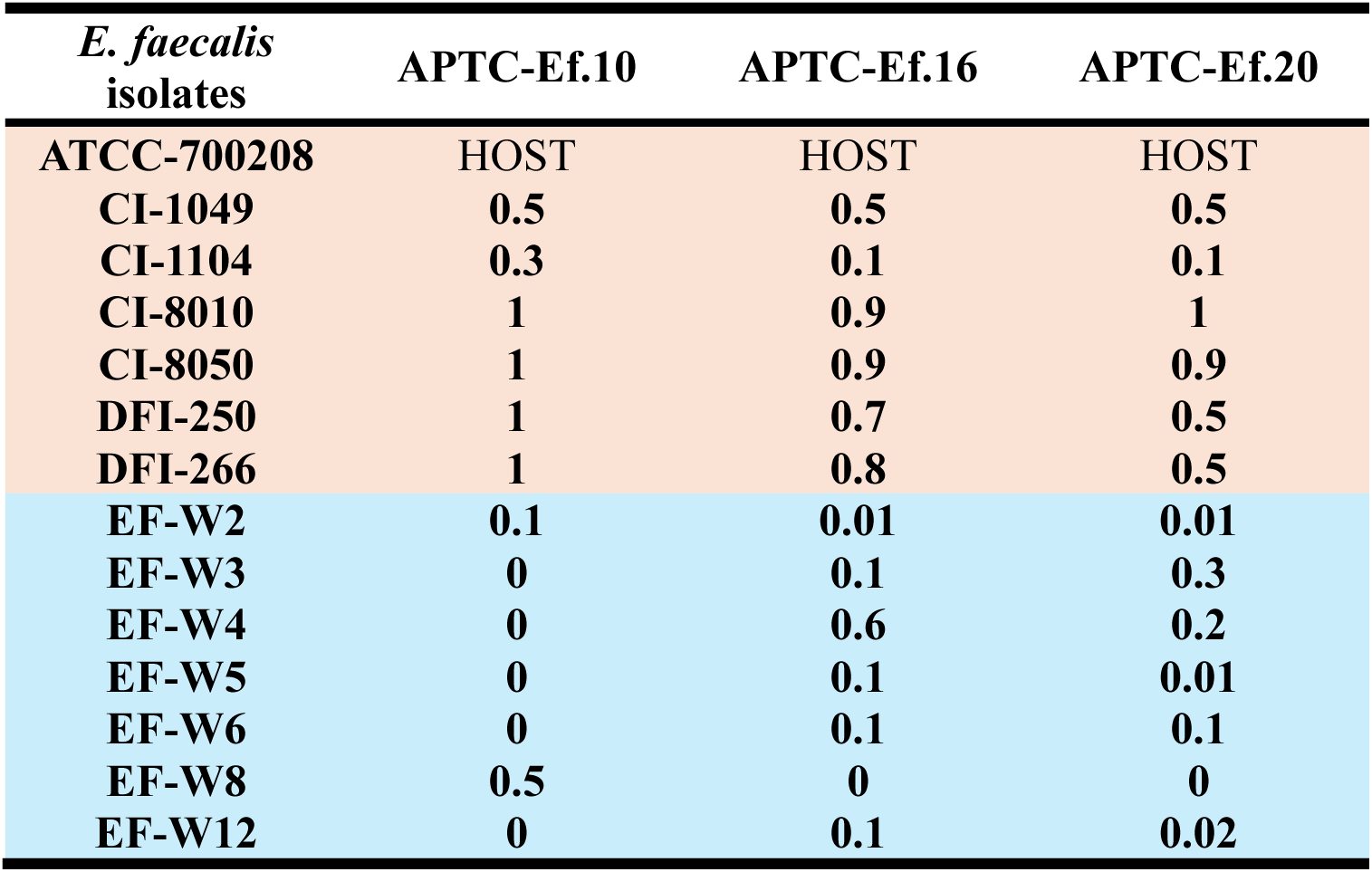
Efficiency of plating of bacteriophages APTC-Ef.10, APTC-Ef.16 and APTC-Ef.20 against *E. faecalis* clinical isolates (highlighted in orange) and wastewater (highlighted in blue). Data presented as the average of 3 biological replicates.

| *E. faecalis*<br>isolates | APTC-Ef.10 | APTC-Ef.16 | APTC-Ef.20 |
| --- | --- | --- | --- |
| ATCC-700208 | HOST | HOST | HOST |
| CI-1049 | 0.5 | 0.5 | 0.5 |
| CI-1104 | 0.3 | 0.1 | 0.1 |
| CI-8010 | 1 | 0.9 | 1 |
| CI-8050 | 1 | 0.9 | 0.9 |
| DFI-250 | 1 | 0.7 | 0.5 |
| DFI-266 | 1 | 0.8 | 0.5 |
| EF-W2 | 0.1 | 0.01 | 0.01 |
| EF-W3 | 0 | 0.1 | 0.3 |
| EF-W4 | 0 | 0.6 | 0.2 |
| EF-W5 | 0 | 0.1 | 0.01 |
| EF-W6 | 0 | 0.1 | 0.1 |
| EF-W8 | 0.5 | 0 | 0 |
| EF-W12 | 0 | 0.1 | 0.02 |

### *E. faecalis* isolates commonly possess a similar linear pseudo temperate bacteriophage and generally have variable genomic and plasmid content correlating with bacteriophage susceptibility

To gain insights in the antimicrobial resistance mechanisms of *E. faecalis* isolates and empirical differences in host range, we generated complete closed genomes with chromosomes and plasmids for all thirteen clinical and wastewater bacterial isolates. Chromosome length varied from 2.60Mbp up to 3.04Mbp. A total of 21 distinct extrachromosomal replicons were carried by these 13 isolates ranging in size from 5.1kbp to 74.8 kbp. Interestingly, five of these were genomically similar linear replicons between 29.4Kbp and 32.1Kbp in size. Bakta and Pharokka annotations of these replicons revealed that these were very closely related (mash distances < 0.01) to the linear pseudo temperate bacteriophage EF62φ that has previously been described to replicate extra-chromosomally as a linear DNA molecule via its encoded *repB* gene in *E. faecalis* [66]. This phage was present in isolates from the various sources, including patients with chronic rhinosinusitis (CI-1104), diabetic foot infection (DFI-250, DFI-266 and CI-8010) and wastewater (EF-W2), though more commonly present in clinically sourced isolates (4/6) compared to wastewater (1/7) (**Table S5**).

Overall, extrachromosomal replicon numbers per isolate ranged from 0 to 5. Notably, 5/7 wastewater isolates did not host any extrachromosomal replicons with the other 2/7 having only 3 combined. In contrast, all 6 clinical isolates contained at least one extrachromosomal replicon, for a total of 18 (**Figure 4**). Of the 16 plasmids (excluding bacteriophage EF62φ), 14 out of 16 were predicted to be mobilizable, while the other 2 were predicted to be non-mobilizable. Of the 2 non-mobilizable plasmids, one (CI-1049 plasmid00002) was in CI-1049, which contained other mobilizable plasmids, while the other (EF-W12 plasmid00001) was the only plasmid present in EF-W12. There was diversity in replication types in the mobilizable plasmids, with the 13 plasmids sharing 9 different replication types. All circular plasmids had close (under 0.1 mash distance to the closest hit) hits in the Mob Typer database.

**Figure 4.**
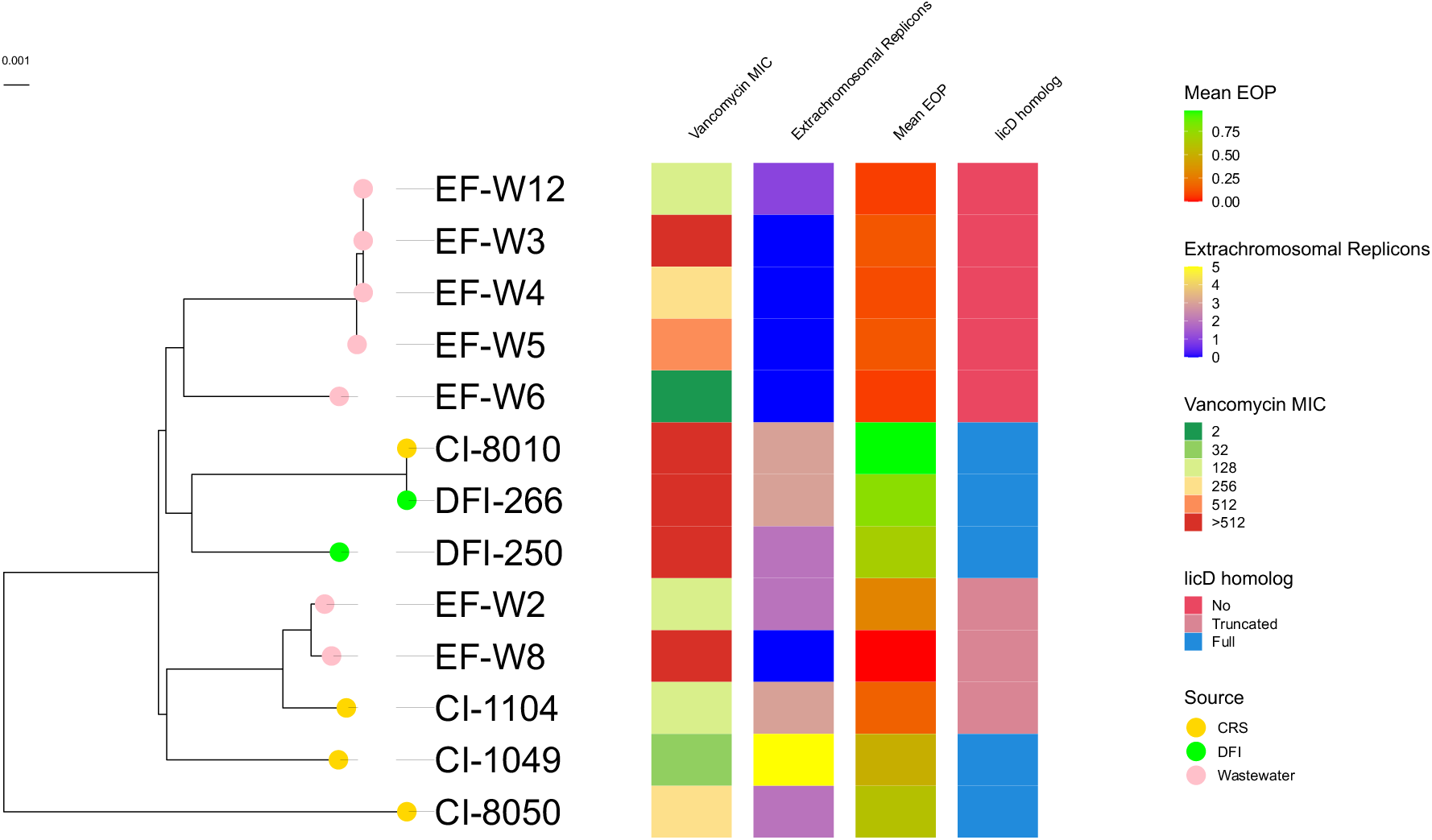
Phylogenetic tree elaborating the genomic similarity between *E. faecalis* isolated from different sources, occurrence of extrachromosomal replicons, vancomycin MIC, and the presence of licD. All wastewater isolates were genetically similar except for EF-W2 and EF-W8 that were closely related to the clinically isolated samples.

In addition to the multiple plasmids found in the isolates, the 13 isolates also contained a variety of genomically predicted antibiotic resistance genes against different antibiotics including quinolones, glycopeptides, Lincosamide, trimethoprim, macrolides, tetracyclines and aminoglycosides. These resistance genes ranged between 2 to 8 genes per isolate, which confirms the wide antibiotic resistance arsenal owned by these isolates (**Table S6**). A phylogenetic tree elaborating the genetic similarities between different *E. faecalis* strains isolated from different sources, occurrence of plasmids, vancomycin MIC, and licD homology is shown in (**Figure 4**).

### Time-kill kinetics of *E. faecalis* phages show bacteriophage insensitive mutant (BIM) are generated within 24 hours

To gain insight into the interaction between the Phages and *E. faecalis*, we constructed a time-kill assay with different MOIs (0.01-100). There was a distinctive difference between the periods each phage could control bacterial growth. APTC-Efa.10 controlled the growth of *E. faecalis* for the first 7 hours, APTC-Efa.16 for the first 12 hours and APTC-Efa.20 for the first 15 hours at all MOIs tested (**Figure 5 a**). In addition, there was a prominent difference between the phages in their dose-dependent antibacterial performance after the initial suppression. High doses of APTC-Efa.10 were necessary to suppress bacterial growth with a direct correlation between MOI and antibacterial activity. At the lowest phage dose tested (MOI 0.01), there was a significant increase in bacterial growth compared to control after the 15-hour time-point. On the other hand, APTC-Efa.16 at high MOI of 10 and 100 efficiently suppressed bacterial growth up to 16 hours post inoculation, after which bacterial growth increased rapidly resulting in a doubling of bacterial cells at the 24-hour timepoint compared to control (p<0.05). However, MOI 1 was the most efficient dose for APTC-Efa.16 and consistently reduced bacterial growth compared to control group after 24 hours. Similarly, APTC-Efa.20 efficiently controlled the bacterial growth at MOI 0.1, however higher phage doses of MOI > 1 resulted in an increase in bacterial growth after 15 hours.

**Figure 5.**
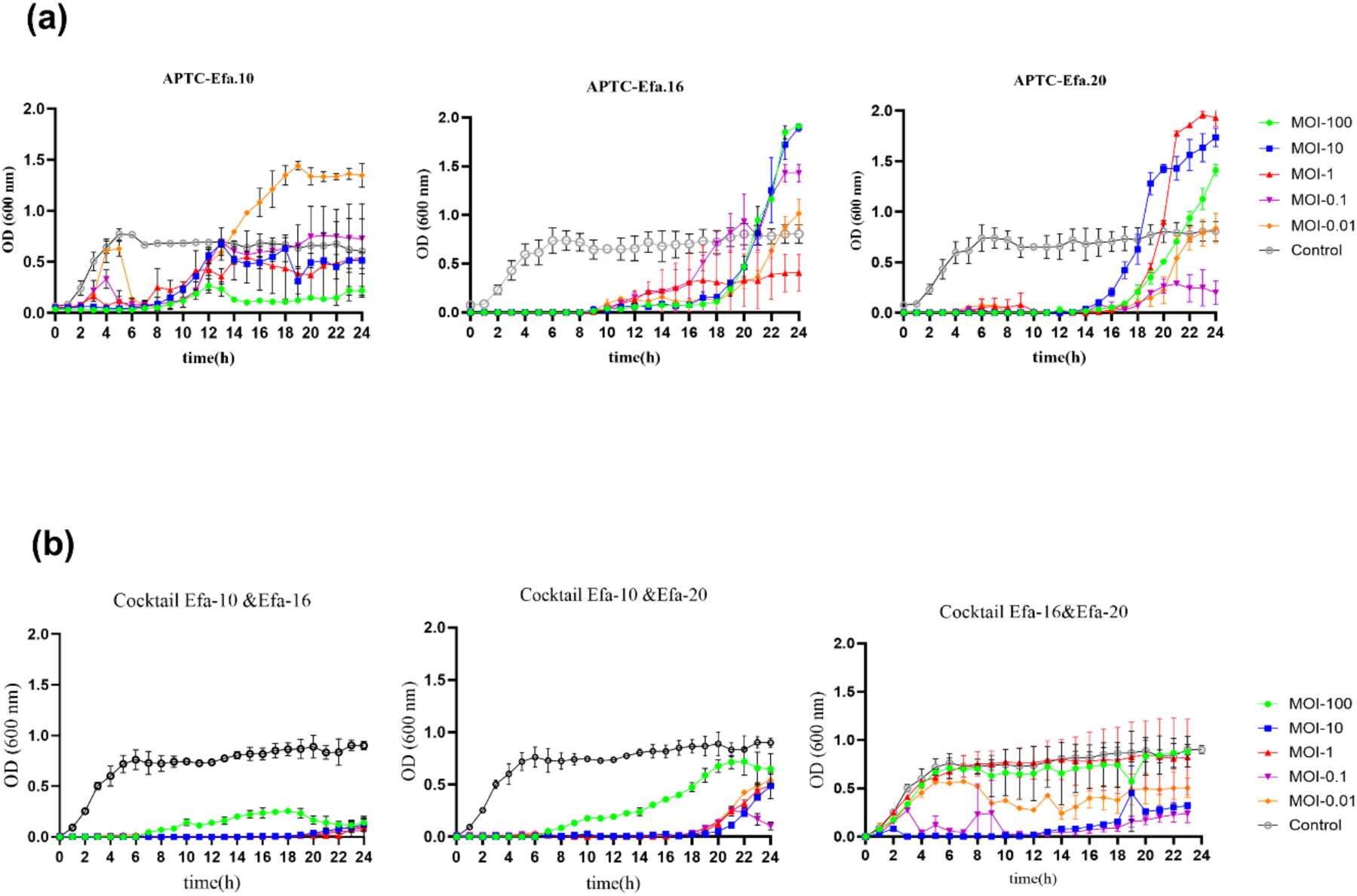
Time-kill kinetics of *Enterococcus* bacteriophages. (a) Growth pattern of *E. faecalis* ATTC 700802 incubated with individual phages APTC-Efa.10, Efa.16 and Efa.20 (MOI) in the range between (0.01-100). (b) Growth pattern of *E. faecalis* ATTC 700802 incubated with 1:1 phage cocktail at different MOIs (0.01-100).

We then prepared a 1:1 cocktail of the phages to determine their potential synergistic antibacterial activity and the time-kill assay was repeated. A cocktail of APTC-Efa.10& APTC-Efa.16 was the most efficient in controlling bacterial growth over 20 hours incubation period at MOIs ranging between 10-0.01 followed by a cocktail of APTC-Efa.10& APTC-Efa.20, which controlled the infection up to 18 hours at the same MOI range. Compared to single phage infections, the onset of BIM formation was significantly delayed when treated with both cocktails. However, MOI 100 was less active and controlled bacterial growth only for six hours in both cocktails. APTC-Efa.16& APTC-Efa.20 was the weakest cocktail as it failed to inhibit bacterial growth better than individual phages in the cocktail. It is noteworthy that lower doses of phage cocktails (MOI 10-0.1) performed better compared to (MOI 100) in all three cocktails (P<0.0001) **(Figure 5 b**). After 24 hours of challenging *E. faecalis* with each of the phages or their cocktails, the surviving bacteria were isolated and their susceptibility to the phages were determined. None of the isolated bacteria were sensitive to the phages and were deemed to be bacteriophage insensitive mutants (BIMs). These included BIM-10, BIM-16 and BIM-20, generated in the presence of APTC-Ef.10, Ef.16 and Ef.20 respectively. DLA assays showed each of those BIMs were resistant to all 3 phages (data not shown).

### Bacteriophage insensitive mutants are less virulent compared to parent strain

The virulence of the BIMs was determined and compared to the parent strain. Firstly, the susceptibility to vancomycin and ampicillin was evaluated. All BIMs were more sensitive to vancomycin and ampicillin scoring MIC values of 8 µg/mL and 0.125 µg/mL respectively compared to 32 µg/mL and 0.5 µg/mL for the parent strain (4-fold reduction in MIC values for BIMs compared to parent strain for both antibiotics).

Next, we tested the capacity of BIMs to form biofilms compared to the parent strain. All BIMs significantly formed lower biofilm biomass compared to the parent strain *P*< 0.0001 (**Figure 6 a)**. Moreover, among the 3 BIMs isolated after incubation with individual phage, BIM-20 showed the lowest biofilm biomass which was significantly lower compared to the other two BIMs. We then evaluated and compared the capacity of the BIMs and parent strain to evade phagocytosis based on their survival within macrophages [64, 65]. There was a significant reduction in average CFUs of *E. faecalis* ATCC-700802 (Log reduction=0.47 and 0.17) in cell lysate compared to BIMs 10 and 20 cell lysates, respectively which indicates a decrease in the ability of THP-1 cells to phagocytose ATCC-700802 compared to BIMs 10 and 20 (**Figure 6b**). However, there was no significant difference in CFU counts between bacteria incubated in media and total bacteria harvested from supernatants and cell lysates for any of the isolates tested (p>0.05) (**Figure S4**) indicating survival of both BIMs and parent strains after phagocytosis.

**Figure 6.**
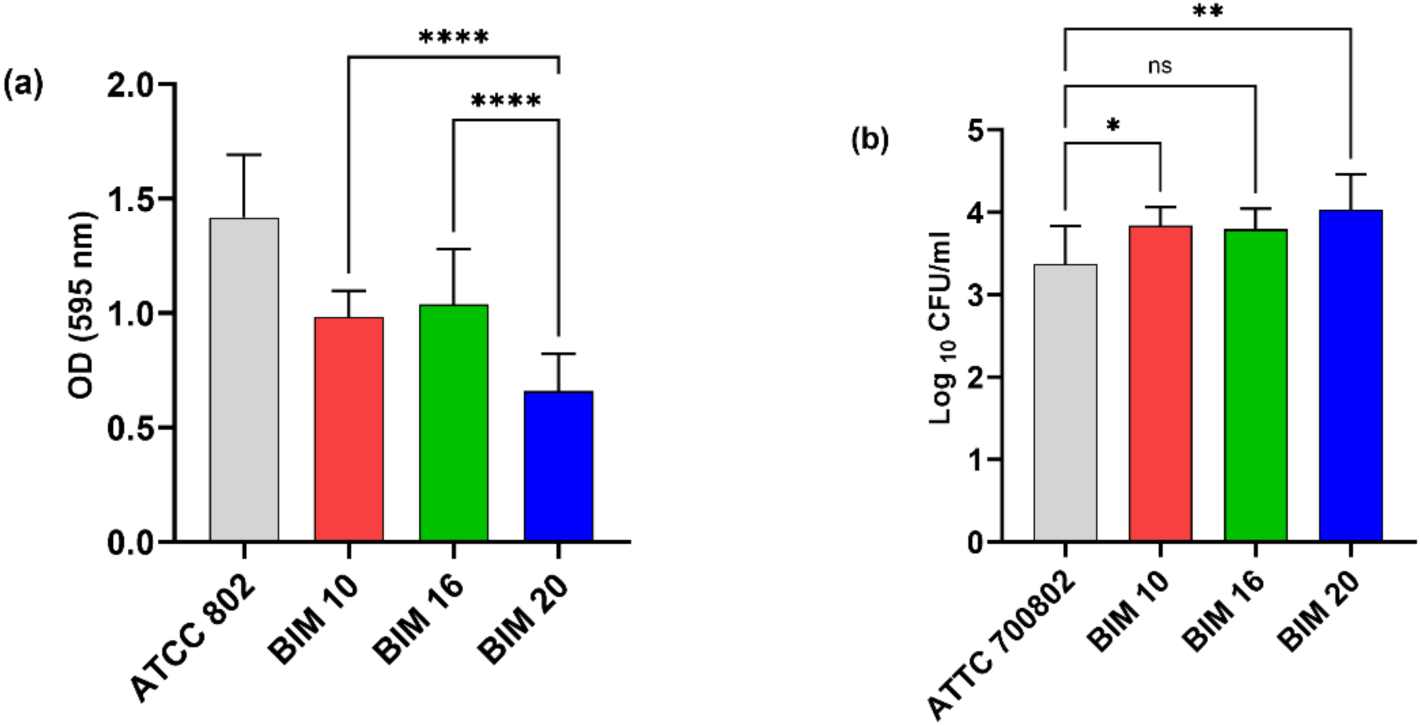
*Assessing virulence capabilities of E. faecalis BIMs following phage infection.* (a) biofilm formation of different BIMs compared to parent ATCC strain. The parent strain formed significantly thicker biofilms compared to BIMs with thinner biofilm was formed by BIM-20 compared to BIM 10 and BIM 16. (b) *E. faecalis* bacteria concentration presented as (log_10_ CFU/mL) in THP-1 cell lysates after incubation at MOI 50 for 1 hour. Bacterial BIMs were phagocytosed more by THP-1 macrophages compared to parent strain with BIM-20 identified as the most vulnerable for phagocytosis. Data presented as mean ± SD of three biological replicates. ns: non-significant, *P* > 0.05, \**P* < 0.05; \*\**P* < 0.01, \*\*\*\**P* < 0.0001 one-way ANOVA test followed by multiple comparisons Dunnett’s test.

Furthermore, we compared the BIM genomes vs the parent ATCC strain to elucidate the genomic basis for the loss of phage susceptibility and attenuated virulence. This indicated that the phosphorylcholine metabolism pathway appeared to be disrupted. All BIMs had variants within or upstream of the phosphorylcholine transferase protein *licD* which is responsible for the transfer of phosphoryl choline to the cell wall glycans such as teichoic acid and lipopolysaccharides [67] (**Table 2**). Specifically, BIM10 had a large structural deletion of 3624 bp, wholly removing *licD* and partially removing a lysozyme encoding gene *LysM* along with the *wcaJ sugar transferase* involved in the biosynthesis of colanic acid, an exopolysaccharide found in bacterial cell wall [68]. (**Figure S5**). All the five other BIMs possessed a small nucleotide variant (SNV) in an intergenic region 109 bp upstream of the 5’-end of the *licD* gene (**supplementary Figures 6-9,11**), while two BIMs that emerged after cocktail treatment APTC-Efa-16&10 and APTC-Efa-20&10 possessed a further SNV 154 bases upstream of the 5’-end of the *licD* gene (**supplementary Figures 7-8**). In addition to the variation in the *licD* locus, BIM 20 had SNVs in 3-hydroxyacyl-ACP dehydratase *FabZ*, and the Nucleoside-diphosphate-sugar epimerase *wcaG* and in the *23S ribosomal RNA* (Supp Figures 9-16), with the *FabZ* mutation being a missense variant (missense_variant c.421T>C p.Ile141Try) and the *wcaG* variant synonymous (synonymous_variant c.508T>A p.Ala169Ala). The *FabZ* variant occurs at the C-terminal end of the protein monomer, specifically at the end of a six-stranded anti-parallel β-sheet with topology 1/2/4/5/6/3 [69] (**Figure 7)**. One BIM isolated after treatment with cocktail composed of APTC-Ef-16 and APTC-Ef-20 was genomically identical to BIM-20 (**Table 2**) (**Supplementary** Figures 16-20).

**Figure 7.**
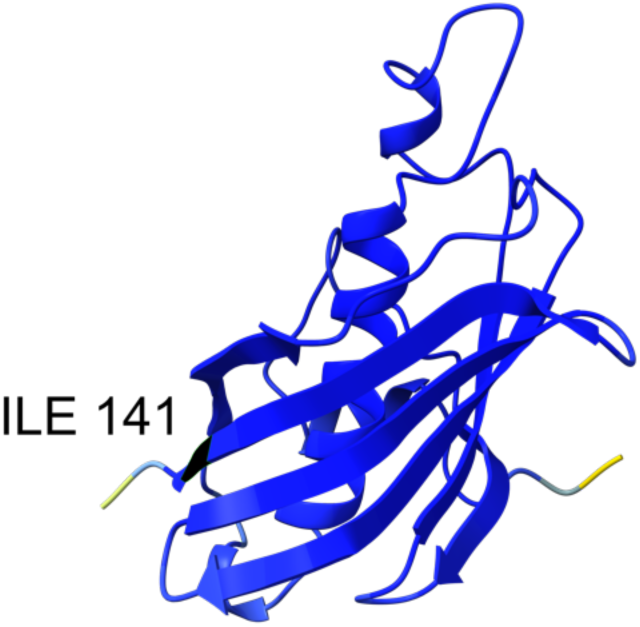
AlphaFold2 predicted structure of the 3-hydroxyacyl-ACP dehydratase FabZ (AFDB accession AF-Q820V3-F1-v4) coloured by predicted local distance difference test (pLDDT), blue indicating residues with pLDDT above 90. The p.Ile141Try variant found in BIM20 at the C-terminal end is highlighted in black.

**Table 2.**
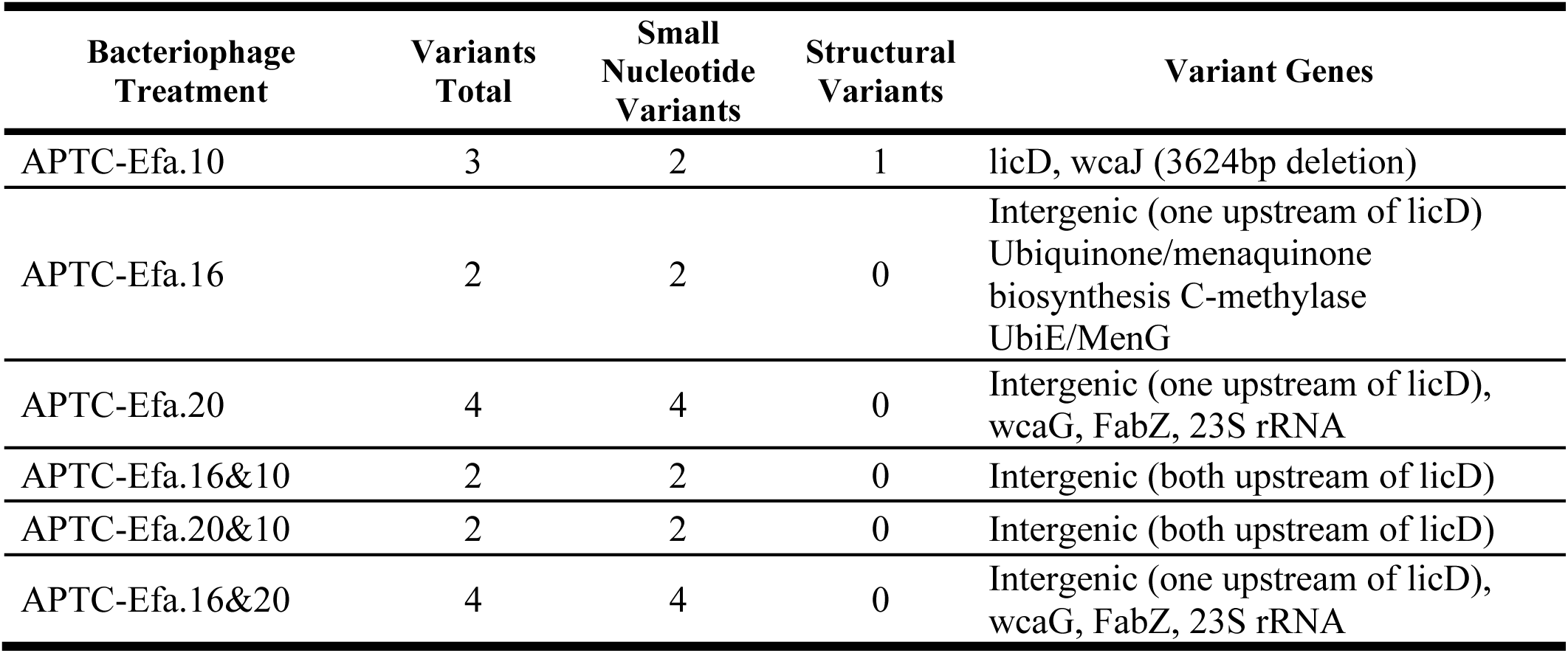
Variants between bacteriophage insensitive mutants and the reference strains.

| Bacteriophage Treatment | Variants Total | Small Nucleotide Variants | Structural Variants | Variant Genes |
| --- | --- | --- | --- | --- |
| APTC-Efa.10 | 3 | 2 | 1 | licD, wcaJ (3624bp deletion) |
| APTC-Efa.16 | 2 | 2 | 0 | Intergenic (one upstream of licD)<br>Ubiquinone/menaquinone biosynthesis C-methylase UbiE/MenG |
| APTC-Efa.20 | 4 | 4 | 0 | Intergenic (one upstream of licD), wcaG, FabZ, 23S rRNA |
| APTC-Efa.16&10 | 2 | 2 | 0 | Intergenic (both upstream of licD) |
| APTC-Efa.20&10 | 2 | 2 | 0 | Intergenic (both upstream of licD) |
| APTC-Efa.16&20 | 4 | 4 | 0 | Intergenic (one upstream of licD), wcaG, FabZ, 23S rRNA |

## Discussion

In this study, we report the isolation and characterization of three lytic phages against *E. faecalis*. Although the three phages are of the same *Kochikohdavirus* genus, they showed variations in their host range and antibacterial activity against a panel of *E. faecalis* strains. APTC-Efa.16 and APTC-Efa.20 had the widest host range, infecting 12 out of 13 tested isolates, while APTC-Efa.10 could only infect 8/13 isolates mostly from samples isolated from infected humans. Moreover, APTC-Efa.16 and APTC-Efa.20 showed higher adsorption rate, burst size and, shorter latent period compared to APTC-Efa.10 suggesting their better control of bacterial infection in clinical settings [70]. Phage cocktails containing APTC-Efa.10 and APTC-Efa.16 or APTC-Efa.20 could efficiently control bacterial growth at various MOIs and could delay the emergence of bacteriophage insensitive mutants (BIMs) compared to single phage infection. Genomic analysis of BIMs indicates a prominent role of the *LicD* gene in conferring phage resistance indicating the potential role of phosphorylcholine as mediator of phage adsorption.

Genomic analysis of the generated BIMs revealed insights regarding the potential target bacterial receptors used by each phage during the infection cycle. Our data showed that the 3 phages likely use the same receptor or different conformations of that receptor to adsorb and initiate the infection, supported by the finding that BIMs generated in the presence of each of the single phages were resistant to all 3 phages. BIM-10, which emerged after infection with APTC-Efa.10, showed a structural variant with the deletion of *LicD* and *WcaJ*, involved in the modulation of the bacterial cell wall [67, 71], while both other BIMs had at least one identical SNV upstream of *LicD*. Whilst experiments are required to investigate this further, we postulate that this might result in a negative regulatory effect of these SNVs on the expression of *LicD* in the BIMs [72, 73]. Together, these findings indicate the dysfunction of the *LicD* gene as important in conferring phage resistance of *E. faecalis* BIMs and that phosphorylcholine can help mediate phage adsorption as shown for *S. pneumoniae* [74]. The *LicD* gene is responsible for the production of the LicD enzyme, a choline phosphotransferase, that transfers phosphorylcholine from bacterial cytoplasm to lipoteichoic acid (LTA) and wall teichoic acid (WTA) on the bacterial surface [75]. The decoration of the bacterial surface with phosphorylcholine has been reported to play crucial rules in pathogenesis of different bacterial species including *S. pneumoniae* and *H. influenzae* allowing the bacteria to modulate host immune responses, colonization of epithelial cells and biofilm formation [67]. To the best of our knowledge the modification of *E. faecalis* cell wall with phosphorylcholine has not yet been reported. However, our data shows that all clinical isolates from humans contained the *LicD* gene, while this gene was absent in the wastewater isolates except for W2 and W8 that had a truncated version of the gene **Figure 4**. This may explain the host range variation of APTC-Efa.10, where the phage could not infect any of wastewater isolates that did not have the *LicD* gene **Table 1**. On the other hand, APTC-Efa.16 and APTC-Efa.20 could infect wastewater isolates that lacked the *LicD* gene but with far lower efficiency compared to clinical isolates harbouring *LicD* indicating the importance of *LicD* and thus phosphorylcholine expression on the bacterial surface for a successful *E. faecalis* phage infection. Our data also suggests possessing *LicD* is evolutionarily favourable for *E. faecalis* isolates that colonise human sinus and diabetic foot niches but is not required and lost in the wastewater environment. Similarly, wastewater isolates mostly lacked mobilizable plasmids, suggesting the traits encoded genomically are not useful in that environment, whereas this is not the case for human-derived *E. faecalis* [76]. Interestingly, this also seemed to be the case with the pseudo-temperate linear bacteriophage found in 4/6 human derived isolates, but only in 1/7 wastewater samples. Previous studies into this phage have indeed suggested it is commonly carried in *E. faecalis* from a variety of human clinical sources [66]. As bacterial genomes adapt to their environment, affecting phage susceptibility, we suggest more diverse collections of isolates be used for phage isolation and host range analyses [77]. Using more diverse hosts for isolation will expand the variety of phages that may be isolated, while using diversely sampled isolates for host-range may help to elucidate possible mechanisms of phage-host interaction, as shown by the analysis of *LicD* in this study.

The other potential target used by APTC-Efa.20 for infecting *E. faecalis* is the lipopolysaccharide capsule. Among the 4 SNVs identified in BIM-20, two were found in *FabZ* and *wcaG*, which are key components for the biosynthesis of unsaturated fatty acids that make up the lipopolysaccharide capsule protecting the bacterial cell wall [68, 69]. Specifically, the missense _ c.421T>C p.Ile141Try *FabZ* SNV at the C-terminal end is likely to affect its termination efficiency and degradation rate [78], while the synonymous variant c.508T>A p.Ala169Ala SNV of *wcaG* may confer fitness effects despite being synonymous [79]. The *E. faecalis* capsule has been reported to increase pathogenicity by promoting immune evasion through the masking of LTA, reducing phagocytosis by macrophages [80, 81]. Furthermore, the capsule contributes to biofilm formation of *E. faecalis* [82]. Our study has demonstrated that BIM-20 produced the lowest biofilm biomass and was more easily phagocytosed by macrophages compared to the parent strain and the other two single-phage BIMs **Figure 6**.

Together these findings support the notion that APTC-Efa.20 might target structures that are essential to the *E. faecalis* capsule and that *FabZ* and/or *wcaG* might play a role in *E. faecalis* biofilm formation. Whilst *wcaG* is known to be important in biofilm formation in *Klebsiella pneumoniae,* the role of this gene in *E. faecalis* has not been specifically explored and warrants investigation [83].

The efficiency by which each phage targets *E. faecalis* could be ascribed to the distinctive differences in the baseplate and tail fibre proteins among the three phages (**Figure 2**). These proteins are responsible for receptor binding on the bacterial surface and the production of lytic enzymes responsible for cell wall disruption such as depolymerases [13]. While depolymerases are commonly produced by phages targeting Gram-negative bacteria, they have also been associated with some previously reported *E. faecalis* phages [84]. Among the 3 phages, the predicted depolymerase domains showed high structural similarity in the beta-propeller depolymerase domain with differences in the lengths of those domains between the phages, suggesting differential attachment to the tail fiber. As tail fiber variation is thought to account for differences in host range and depolymerase activities between phages [77, 85–87], the genomic variation in this tail fiber protein potentially explains the observed difference in biofilm reduction and efficiency of plating between the 3 phages.

Previous research has shown promise to reduce the occurrence of BIMs by designing phage cocktails where each phage in the cocktail targets a different bacterial receptor [88]. Our data shows that cocktails composed of APTC-Efa.10 with either APTC-Efa.16 or APTC-Efa.20 delayed the occurrence of BIMs and had better antibacterial performance compared to single phages. In contrast, cocktails composed of APTC-Efa.16 and APTC-Efa.20 had worse antibacterial performance compared to single phage treatments. These findings might be caused by variable host-phage interactions resulting from differences in tail fibres among the phages. Whilst APTC-Efa.16 &20 harbour depolymerase domains that are very similar at both sequence and structural levels, APTC-Efa.10 is more divergent at the sequence level (**Figure S2**). As AlphaFold predictions are designed to be static snapshots and do not capture dynamics or conformational diversity well [89, 90], it is plausible that the APTC-Efa.10 depolymerase has different *in vitro* molecular dynamics compared to the other two, despite a similar static structure. Therefore, whilst all 3 phages might target the same receptor on the bacterial surface, they might do that by interacting with different conformations of that receptor resulting in a delay of BIM formation. This research furthermore highlights the importance of testing phage cocktails before their use in the clinical setting [91].

Despite the inability of the phages to completely inhibit bacterial regrowth, the surviving colonies were more vulnerable to antibiotics with 4-fold reduction in MIC values. These BIMs were also less virulent as they formed weaker biofilms and reduced their ability to evade phagocytosis (**Figure 6)**. Furthermore, the strong antibiofilm activity of these phages underpin their suitability to control *E. faecalis* infection, given its notoriety for forming biofilms [84, 92]. Since the reduction of virulence and antimicrobial resistance of phage resistant isolates is a reported phenomenon [93], we suggest that the phages isolated in this study could be combined synergistically with antibiotics in a therapeutic context [94]. Further studies optimizing the combination of phages and antibiotics is crucial to better understand the potential effect especially against biofilms [95].

## Conclusion

Three novel *Kochikohdavirus* bacteriophages targeting *E. faecalis* were isolated and characterized. All three phages showed strong lytic activity against vancomycin resistant *E. faecalis* both in planktonic and biofilms modes of growth. The phages showed better performance against *E. faecalis* human-derived clinical isolates containing the phosphorylcholine transferase gene *LicD* compared to wastewater derived isolates lacking it, suggesting a strong correlation between phosphorylcholine presence in cell wall and phage adsorption to the cells. Furthermore, we demonstrated that the isolated APTC-Efa phages can disrupt biofilms, with variations in host range and biofilm reduction likely caused by variation in tail fiber genes containing a depolymerase domain. Moreover, the isolated BIMs following phage treatment were less virulent and more sensitive to antibiotics, with 4-fold lower MIC values for vancomycin and ampicillin. Future studies combining the isolated phages with different antibiotics are warranted to support their potential clinical application.

## Authors contribution

M.A. writing original draft, methodology, data analysis and visualization; G.B. writing original draft, methodology, data analysis and visualization; S.F. methodology and data visualization; S.R.G. data analysis and visualization; P.JW., A.J.P. and S.V. Supervision, conceptualization, funding acquisition, writing reviewing and editing.

## Data availability

All sequencing reads for phage, bacterial isolates and BIMs can be found in the Sequence Read Archive (SRA) under BioProject PRJNA1260167. All bioinformatics code and other data (including the protein structure predictions for every phage protein) used to reproduce the analyses in this paper can be found at https://github.com/gbouras13/Awad_et_al_LicD.

## Acknowledgements

The authors acknowledge the instruments and scientific and technical assistance of Microscopy Australia at Adelaide Microscopy, The University of Adelaide, a facility that is funded by the University, and State and Federal Governments. This work was supported with supercomputing resources provided by the Phoenix HPC service at the University of Adelaide and on Setonix at the Pawsey Supercomputing Research Centre with funding from the Australian Government and the Government of Western Australia.

