## Supplementary Materials for "LicD-Mediated Cell Wall Decoration Governs Phage Sensitivity in Enterococcus faecalis Clinical Isolates"

<u>**Supplementary data**</u>

### A. Supplementary tables

**Table S 1. Source of bacterial isolates and minimum inhibitory concentrations against ampicillin and vancomycin (n=3) CRSwNP= chronic rhinosinusitis with nasal Polyps patient; DFI= diabetic foot infection patient**

| ***E. faecalis* isolate** | **Source** | **Ampicillin MIC (µg/mL)** | **Vancomycin MIC (µg/mL)** |
|---|---|---|---|
| **ATCC-700802** | ATCC strain | 0.5 | 32 |
| **CI-1049** | CRSwNP | 128 | 32 |
| **CI-1104** | CRSwNP | 256 | 128 |
| **CI-8010** | DFI | >512 | >512 |
| **CI-8050** | DFI | >512 | 256 |
| **DFI-250** | DFI | >512 | >512 |
| **DFI-266** | DFI | >512 | >512 |
| **Efa-W2** | QEH wastewater | 64 | 128 |
| **Efa-W3** | QEH wastewater | 256 | >512 |
| **Efa-W4** | QEH wastewater | > 512 | 256 |
| **Efa-W5** | QEH wastewater | 256 | 512 |
| **Efa-W6** | QEH wastewater | 16 | 2 |
| **Efa-W8** | QEH wastewater | >512 | >512 |
| **Efa-W12** | QEH wastewater | 256 | 128 |

**Table S 2. Bacteriophage host range against isolated E. faecalis strains.**

| **Bacteria isolate** | **Bacterial source** | **APTC-EF10** | **APTC-EF16** | **APTC-EF20** |
|---|---|---|---|---|
| ATCC-700802 | ATCC | HOST | HOST | HOST |
| CI-1049 | ENT patient | + | + | + |
| CI-1104 | ENT patient | + | +/- | +/- |
| DFI-8010 | Diabetic foot wound | + | + | + |
| DFI-8050 | Diabetic foot wound | + | + | + |
| DFI-250 | Diabetic foot wound | + | + | + |
| DFI-266 | Diabetic foot wound | + | + | + |
| EF-W2 | QEH wastewater | - | +/- | +/- |
| EF-W3 | QEH wastewater | - | +/- | +/- |
| EF-W4 | QEH wastewater | - | +/- | +/- |
| EF-W5 | QEH wastewater | - | +/- | +/- |
| EF-W6 | QEH wastewater | - | +/- | +/- |
| EF-W8 | QEH wastewater | - | +/- | +/- |
| EF-W12 | QEH wastewater | - | +/- | +/- |

**Table S 3. List of bacteria used for determining the specificity of bacteriophages.**

| Bacteria isolate | APTC-EF10 | APTC-EF16 | APTC-EF20 |
|---|---|---|---|
| E. faecium ATCC 1559 | - | - | - |
| *S. aureus* RN4220 | - | - | - |
| *S. aureus* CI 216 | - | - | - |
| *S. aureus* CI 226 | - | - | - |
| *S. aureus* CI 256 | - | - | - |
| *S. aureus* CI 047 | - | - | - |
| *S. lugdunensis* CI 1134 | - | - | - |
| *S. lugdunensis* CI 1136 | - | - | - |
| *S. epidermidis* CI 002 | - | - | - |
| *S. epidermidis* CI 1113 | - | - | - |
| *S. epidermidis* CI 1122 | - | - | - |

**Table S 3. Phage genome sequencing and annotation characteristics by Pharokka**

| Phage | Assigned genus | Closest INPHARED match | Length (bp) | GC content (%) | CDS count |
|---|---|---|---|---|---|
| APTC-Ef.10 | Kochikohdavirus | Enterococcus phage vB_EfaM_Ef2.3 | 143456 | 35.83 | 239 |
| APTC-Ef.16 | Kochikohdavirus | Enterococcus phage vB_EfaM_LG1 | 145038 | 35.96 | 244 |
| APTC-Ef.20 | Kochikohdavirus | Enterococcus phage vB_OCPT_Bob | 138438 | 35.76 | 235 |

**Table S 4. Phage genome annotation table by PHROG category**

| Description | Ef-10 | Ef-16 | Ef-20 |
|---|---|---|---|
| **CDS** | 239 | 244 | 235 |
| **connector** | 0 | 0 | 0 |
| **DNA, RNA and nucleotide metabolism** | 30 | 34 | 29 |
| **head and packaging** | 20 | 18 | 20 |
| **integration and excision** | 0 | 0 | 0 |
| **lysis** | 2 | 2 | 2 |
| **moron, auxiliary metabolic gene and host takeover** | 8 | 9 | 8 |
| **other** | 12 | 12 | 11 |
| **tail** | 13 | 13 | 12 |
| **transcription regulation** | 3 | 3 | 3 |
| **unknown function** | 151 | 153 | 150 |
| **VFDB (Virulence Factors)** | 0 | 0 | 0 |
| **CARD (AMR)** | 0 | 0 | 0 |
| **ACR (anit-CRISPR)** | 0 | 0 | 0 |
| **Defensefinder** | 1 | 1 | 1 |
| **Netflax** | 0 | 0 | 0 |

**Table S 5. A comparison between the genomic data of *E. faecalis* isolates from different sources demonstrating the total genome length and plasmid plasmids presence.**

| Isolates | Source | Total Genome Length | Contigs | Chromosome Length | Long Read Coverage | Extrachromosomal replicons | Pseudo temperate linear bacteriophage |
|---|---|---|---|---|---|---|---|
| DFI-250 | DFI | 3131600 | 3 | 3033464 | 76 | 2 | Yes |
| DFI-266 | DFI | 3177527 | 4 | 3014868 | 65 | 3 | Yes |
| CI-1049 | CRSwNP | 3050661 | 6 | 2840277 | 49 | 5 | -- |
| CI-1104 | CRSwNP | 3042631 | 4 | 2900602 | 60 | 3 | Yes |
| CI-8010 | CRSwNP | 3178663 | 4 | 3014988 | 68 | 3 | Yes |
| CI-8050 | CRSwNP | 2682477 | 3 | 2602574 | 93 | 2 | -- |
| EF-W12 | Wastewater | 3069699 | 2 | 3056870 | 73 | 1 | -- |
| EF-W2 | Wastewater | 3005951 | 3 | 2921402 | 57 | 2 | Yes |
| EF-W3 | Wastewater | 3043325 | 1 | 3043325 | 35 | 0 | -- |
| EF-W4 | Wastewater | 3044041 | 1 | 3044041 | 44 | 0 | -- |
| EF-W5 | Wastewater | 3007472 | 1 | 3007472 | 40 | 0 | -- |
| EF-W6 | Wastewater | 2954722 | 1 | 2954722 | 83 | 0 | -- |
| EF-W8 | Wastewater | 3070783 | 3 | 2959622 | 34 | 0 | -- |

**Table S 6. Genomic prediction of antibiotic resistance genes in *E. faecalis* isolated strains.**

| Sample | Total Genes | LINCOSAMIDE/STREPTOGRAMIN | GLYCOPEPTIDE | TETRACYCLINE | OTHER (HEAT) | BACITRACIN | PHENICOL | LINCOSAMIDE/MACROLIDE/STREPTOGRAMIN | QUINOLONE | AMINOGLYCOSIDE | TRIMETHOPRIM |
|---|---|---|---|---|---|---|---|---|---|---|---|
| CI-1049 | 7 | 1 | 0 | 2 | 0 | 4 | 0 | 0 | 0 | 0 | 0 |
| CI-1104 | 4 | 1 | 1 | 1 | 0 | 0 | 0 | 1 | 0 | 0 | 0 |
| CI-8010 | 8 | 1 | 0 | 1 | 0 | 0 | 0 | 2 | 2 | 1 | 1 |
| CI-8050 | 2 | 1 | 0 | 0 | 1 | 0 | 0 | 0 | 0 | 0 | 0 |
| DFI-250 | 3 | 1 | 0 | 1 | 0 | 0 | 1 | 0 | 0 | 0 | 0 |
| DFI-266 | 8 | 1 | 0 | 1 | 0 | 0 | 0 | 2 | 2 | 1 | 1 |
| EF-W12 | 7 | 1 | 0 | 1 | 0 | 0 | 1 | 1 | 2 | 1 | 0 |
| EF-W2 | 3 | 1 | 1 | 1 | 0 | 0 | 0 | 0 | 0 | 0 | 0 |
| EF-W3 | 7 | 1 | 0 | 1 | 0 | 0 | 1 | 1 | 2 | 1 | 0 |
| EF-W4 | 7 | 1 | 0 | 1 | 0 | 0 | 1 | 1 | 2 | 1 | 0 |
| EF-W5 | 7 | 1 | 0 | 1 | 0 | 0 | 1 | 1 | 2 | 1 | 0 |
| EF-W6 | 3 | 1 | 0 | 1 | 1 | 0 | 0 | 0 | 0 | 0 | 0 |
| EF-W8 | 3 | 1 | 1 | 1 | 0 | 0 | 0 | 0 | 0 | 0 | 0 |

### B. Supplementary figures

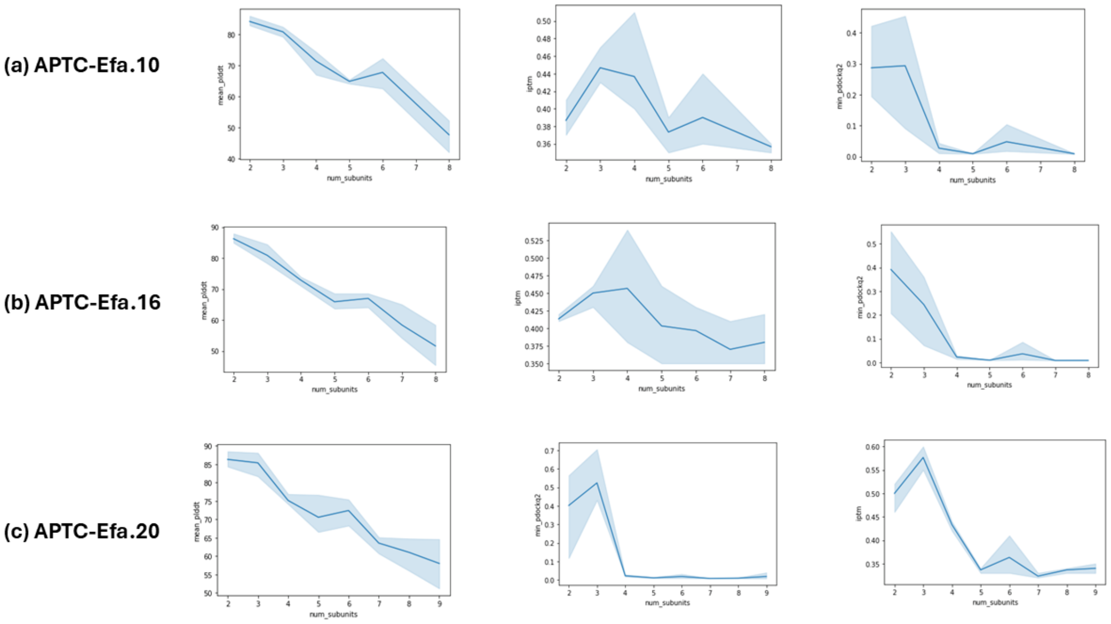


Figure S1. Metrics used for oligomeric state prediction of *APTC-Ef.10, APTC-Ef.16 and APTC-Ef.20*. Mean pLDDT score, ipTM score, and minimum DockQ2 score from three structures generated with 2-9 chain copies. The pLDDT score provides a measure of the quality of the protein structure and the ipTM score and pDockQ2 provide a measure of the extent of interaction between chains.

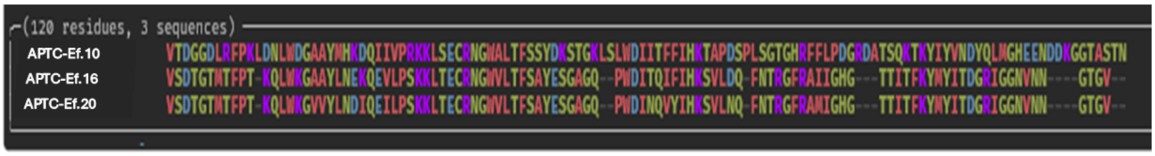


(a)

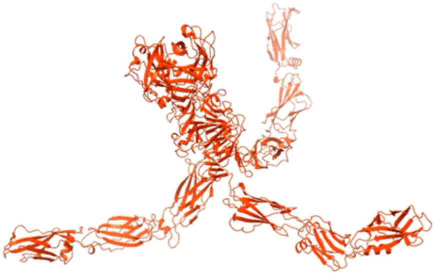

(b)

**Figure S 2.** (a) Predicted beta-propeller depolymerase domain multiple sequence alignment. The predicted depolymerase domain of APTC-EF.20 is aligned to the homologous region of the equivalent APTC-EF.16 and APTC-Ef.10 tail fibre proteins. The homologous region of APTC-EF.16 is highly similar (9 single residue differences), while APTC-Ef.10 is extremely divergent. (b) Trimer of APTC-EF.20: APTC-EF.20_CDS_0029.

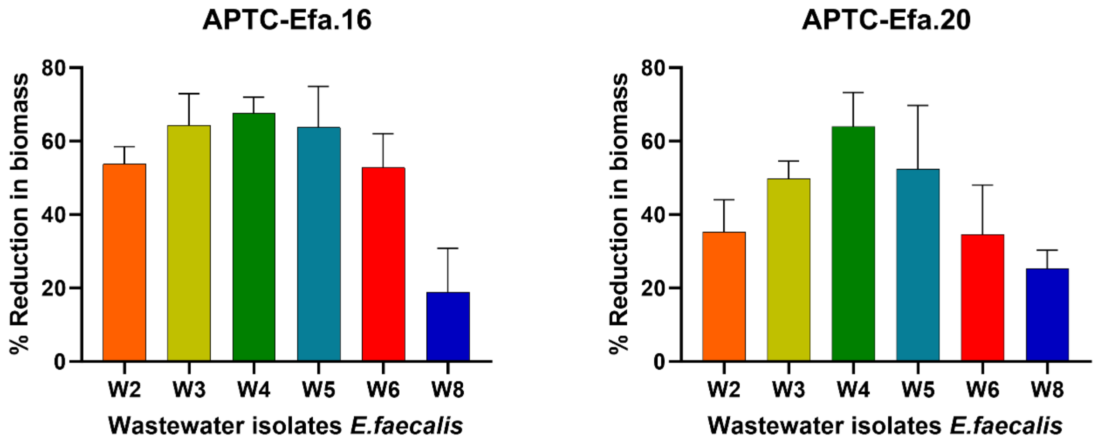


**Figure S 3.** Antibiofilm activity of APTC-Efa.16 and APTC-Efa.20 against E. faecalis isolated from wastewater. Data presented as percent reduction in biofilm biomass compared to untreated control biofilms.

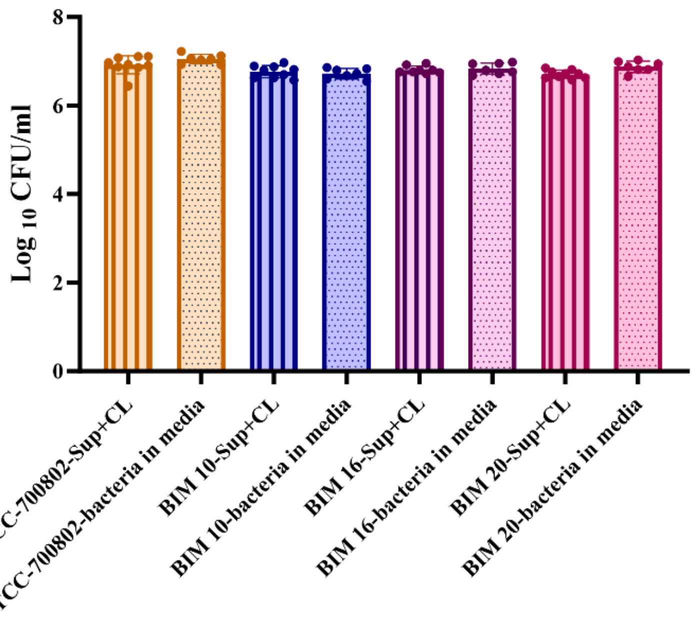


**Figure S 4.** Total colony forming unit (CFU) of supernatant (Sup) and cell lysate (CL) vs bacteria in media (3 biological replicates). Data represent the mean ± SD of three biological replicates, two-way analysis of variance (2-way ANOVA).

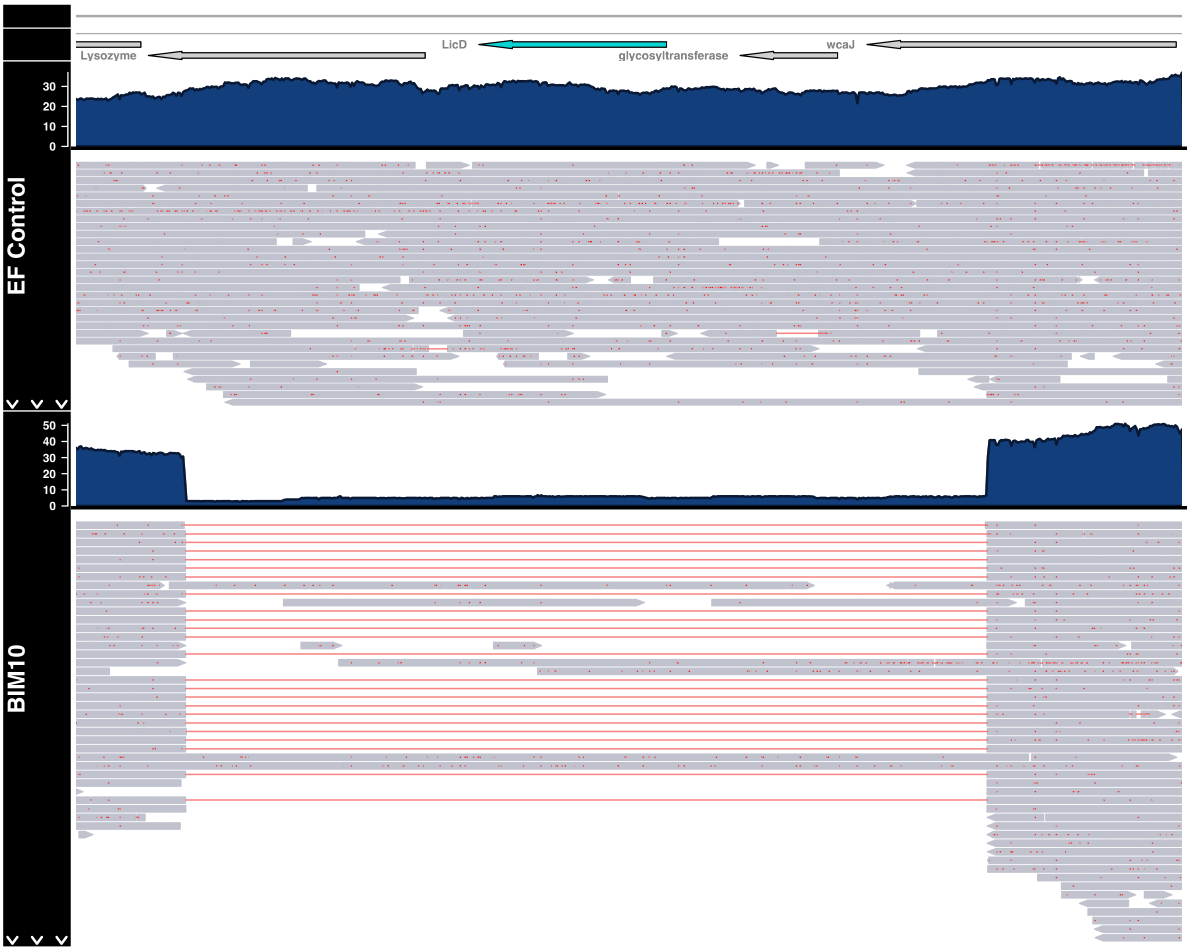


**Figure S 5.** Pileup plot showing the 3624bp structural deletion in BIM-10 compared to the reference genome ATCC-700802. The genomic context is show at the top with the LicD highlighted in blue. In each panel, the read coverage is shown in blue, each mapped read is represented in grey, with red lines indicating a deletion. The control (ATCC-700802) is shown in the top panel, while BIM10 is shown below.

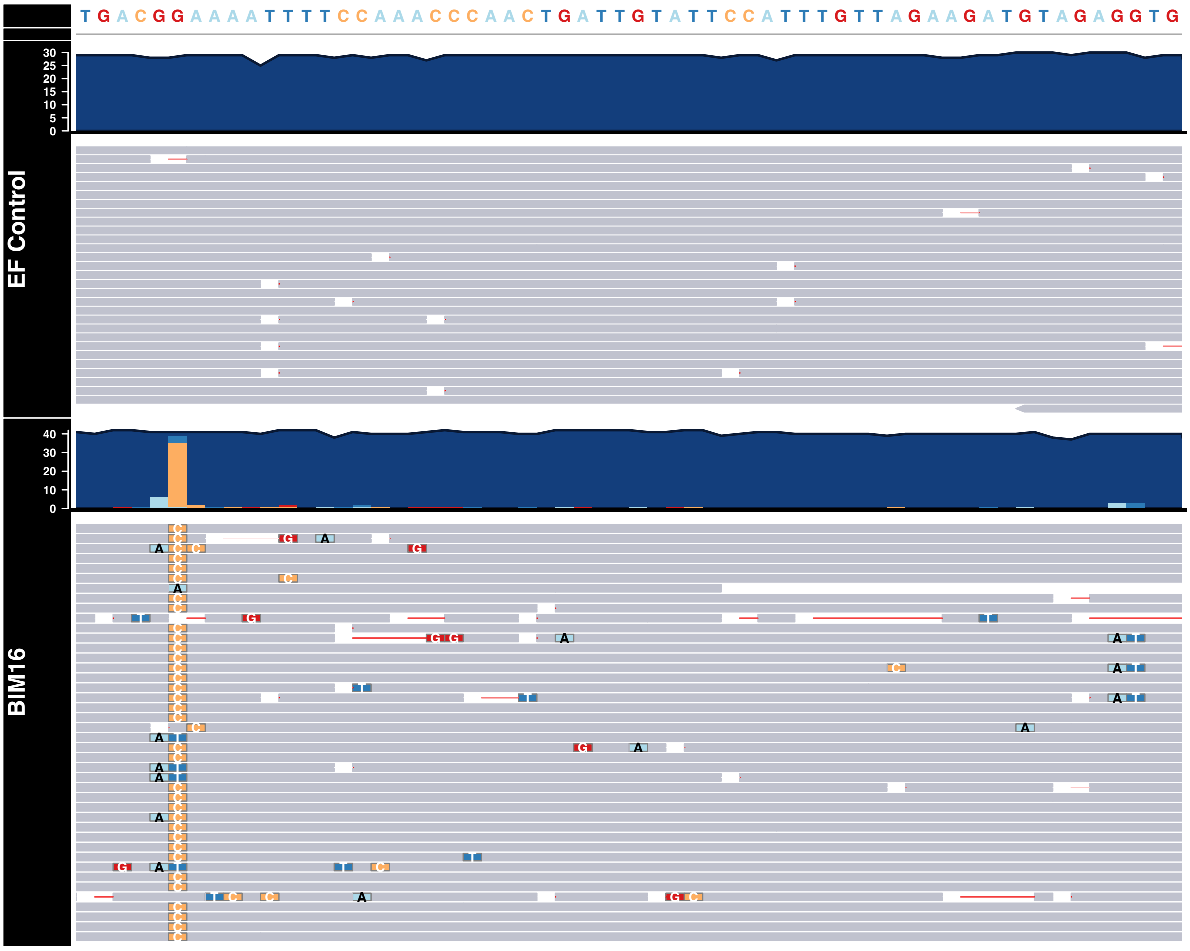


**Figure S 6.** Pileup plot showing the G->C SNV 109 bases upstream of LicD in BIM-16 compared to the reference genome ATCC-700802. In each panel, the read coverage is shown in blue, each mapped read is represented in grey, with yellow indicating a G->C variant. The control (ATCC-700802) is shown in the top panel, while BIM-16 is shown below.

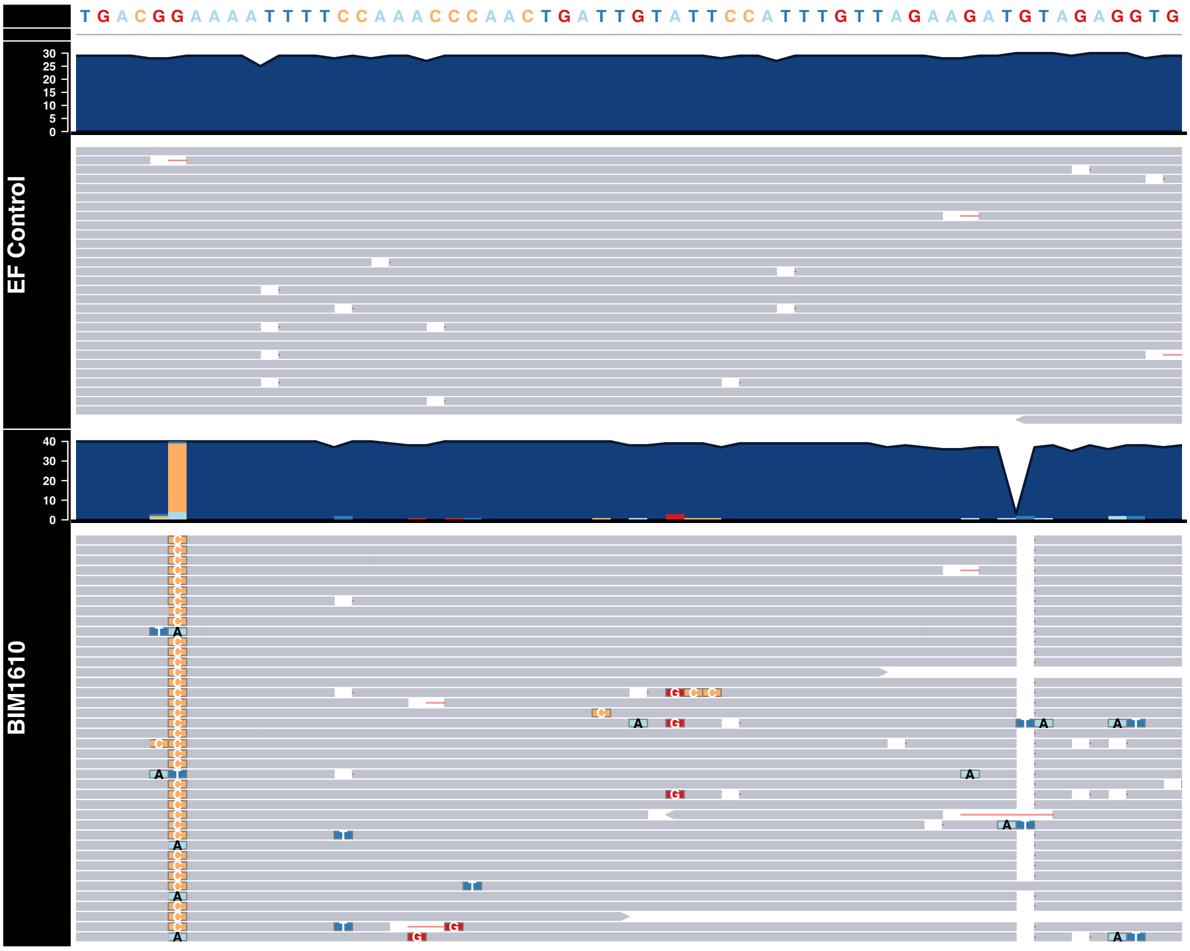


**Figure S 7.** Pileup plot showing the G->C SNV 109 bases upstream of LicD and the TG->T small deletion 154 bases upstream in BIM-1610 compared to the reference genome ATCC-700802. In each panel, the read coverage is shown in blue, each mapped read is represented in grey, with yellow indicating the G->C variant and the gap to the right indicates the TG->T small deletion. The control (ATCC-700802) is shown in the top panel, while BIM-1610 is shown below.

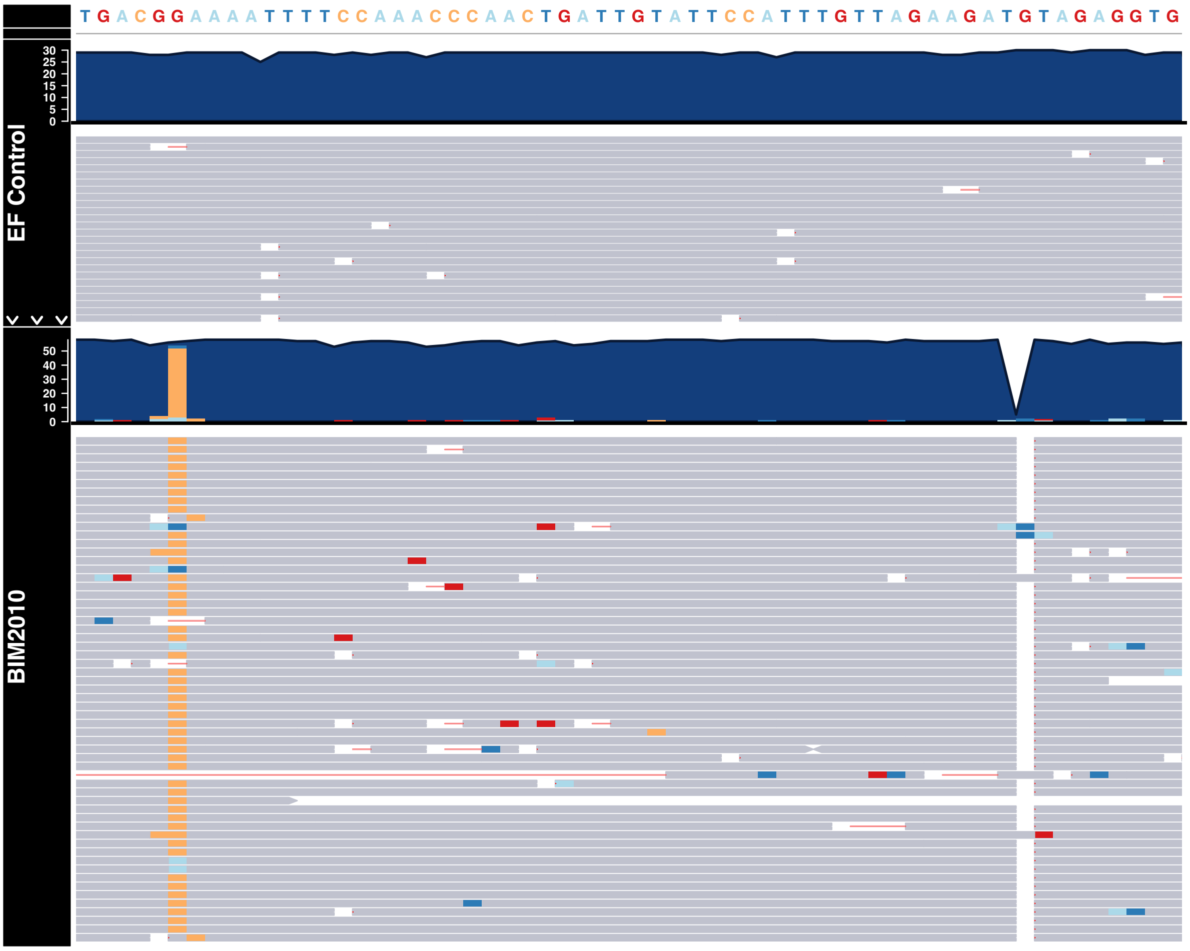


**Figure S 8.** Pileup plot showing the G->C SNV 109 bases upstream of LicD and the TG->T small deletion 154 bases upstream in BIM-2010 compared to the reference genome ATCC-700802. In each panel, the read coverage is shown in blue, each mapped read is represented in grey, with yellow indicating the G->C variant and the gap to the right indicates the TG->T small deletion. The control (ATCC-700802) is shown in the top panel, while BIM-2010 is shown below.

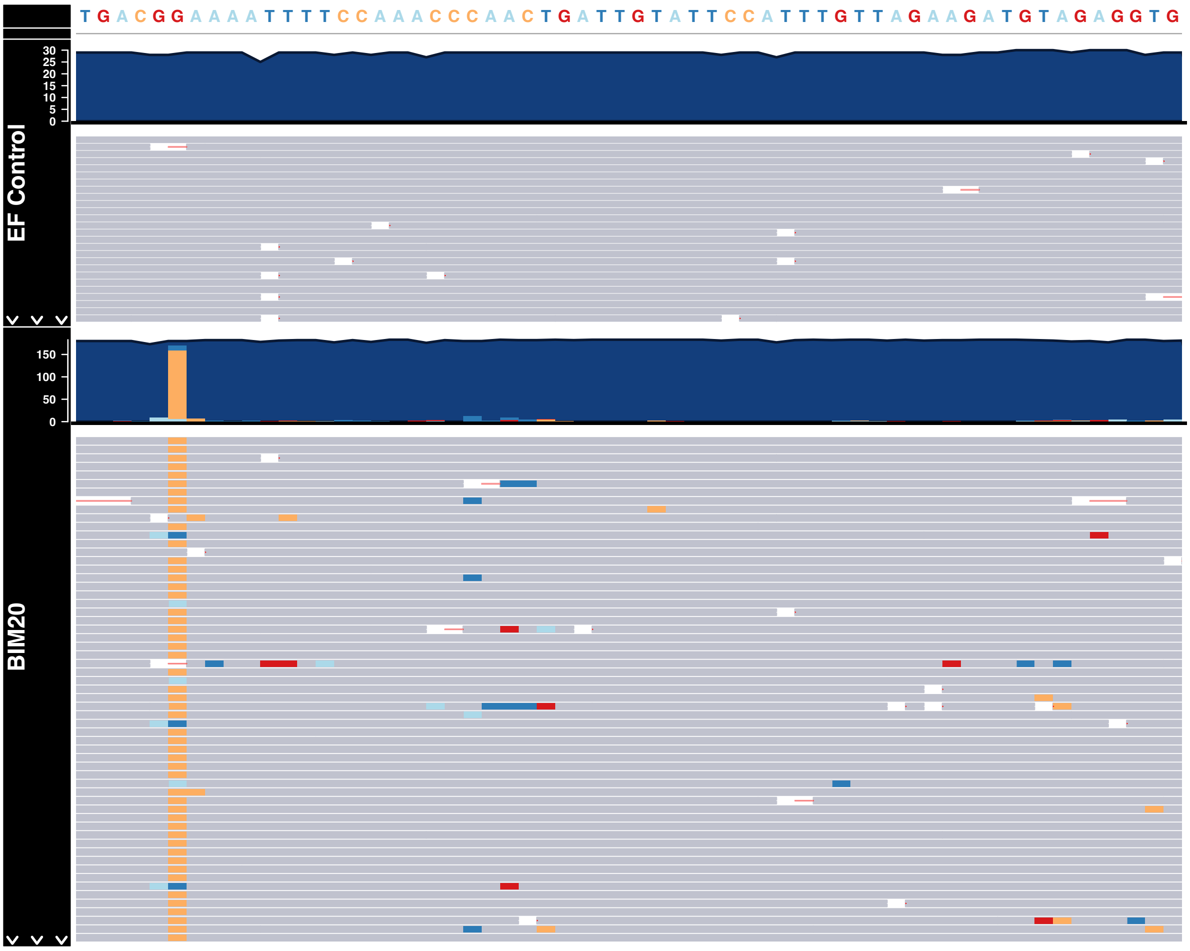


**Figure S 9.** Pileup plot showing the G->C SNV 109 bases upstream of LicD in BIM-20 compared to the reference genome ATCC-700802. In each panel, the read coverage is shown in blue, each mapped read is represented in grey, with yellow indicating the G->C variant. The control (ATCC-700802) is shown in the top panel, while BIM20 is shown below.

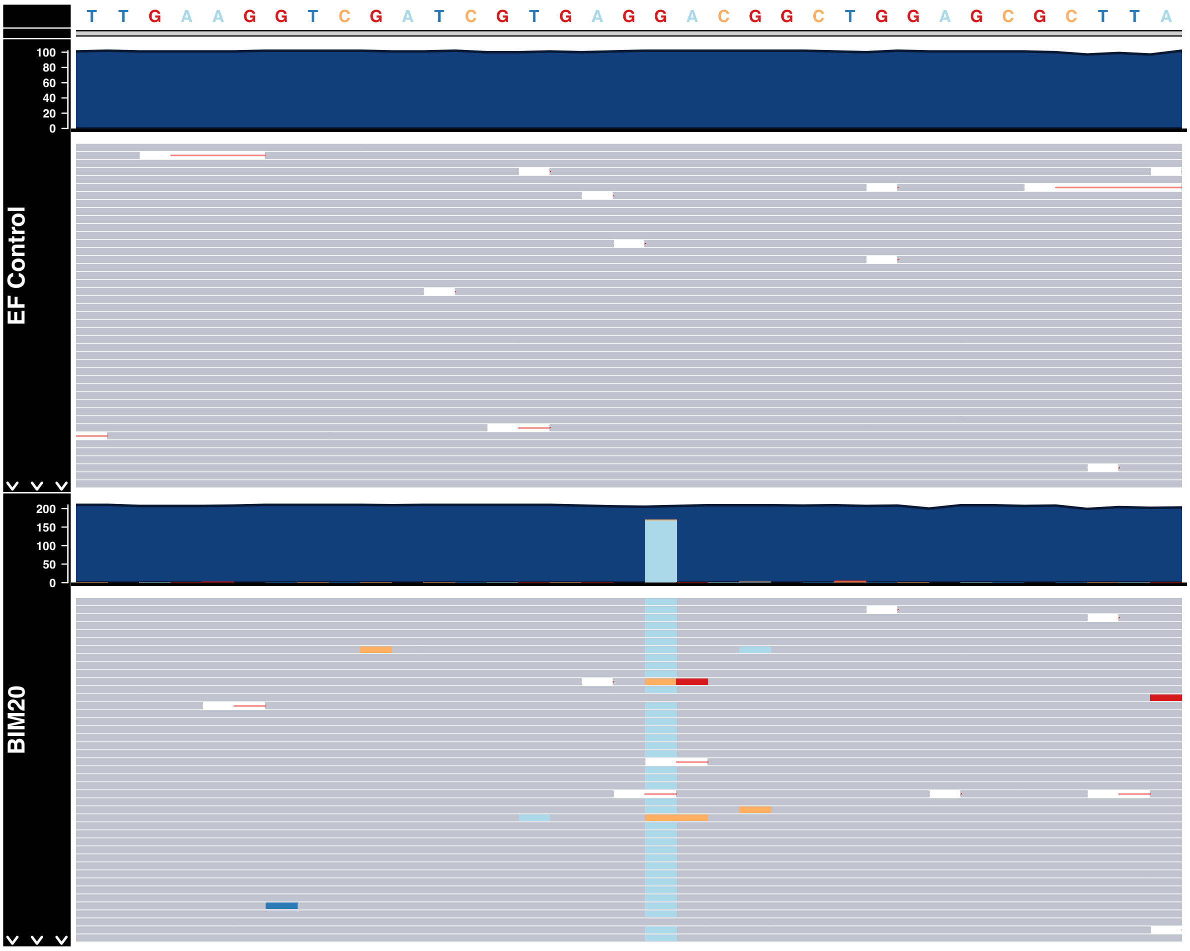


**Figure S 10.** Pileup plot showing the C->A SNV in 23S rRNA gene in BIM-20 compared to the reference genome ATCC-700802. In each panel, the read coverage is shown in blue, each mapped read is represented in grey, with light blue indicating the variant. The control (ATCC-700802) is shown in the top panel, while BIM20 is shown below.

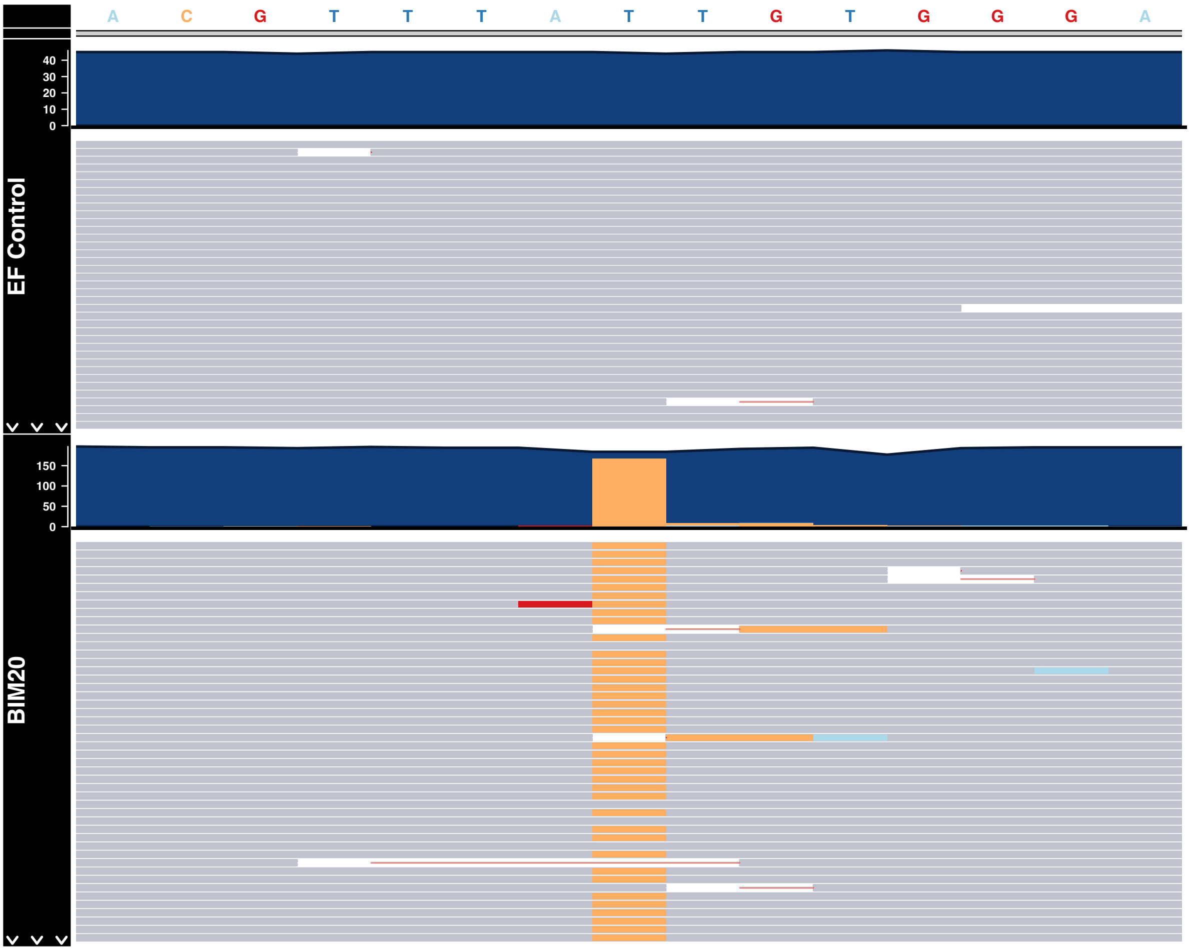


**Figure S 11.** Pileup plot showing the T->C SNV in FabZ gene in BIM-20 compared to the reference genome ATCC-700802. In each panel, the read coverage is shown in blue, each mapped read is represented in grey, with yellow indicating the variant. The control (ATCC-700802) is shown in the top panel, while BIM20 is shown below.

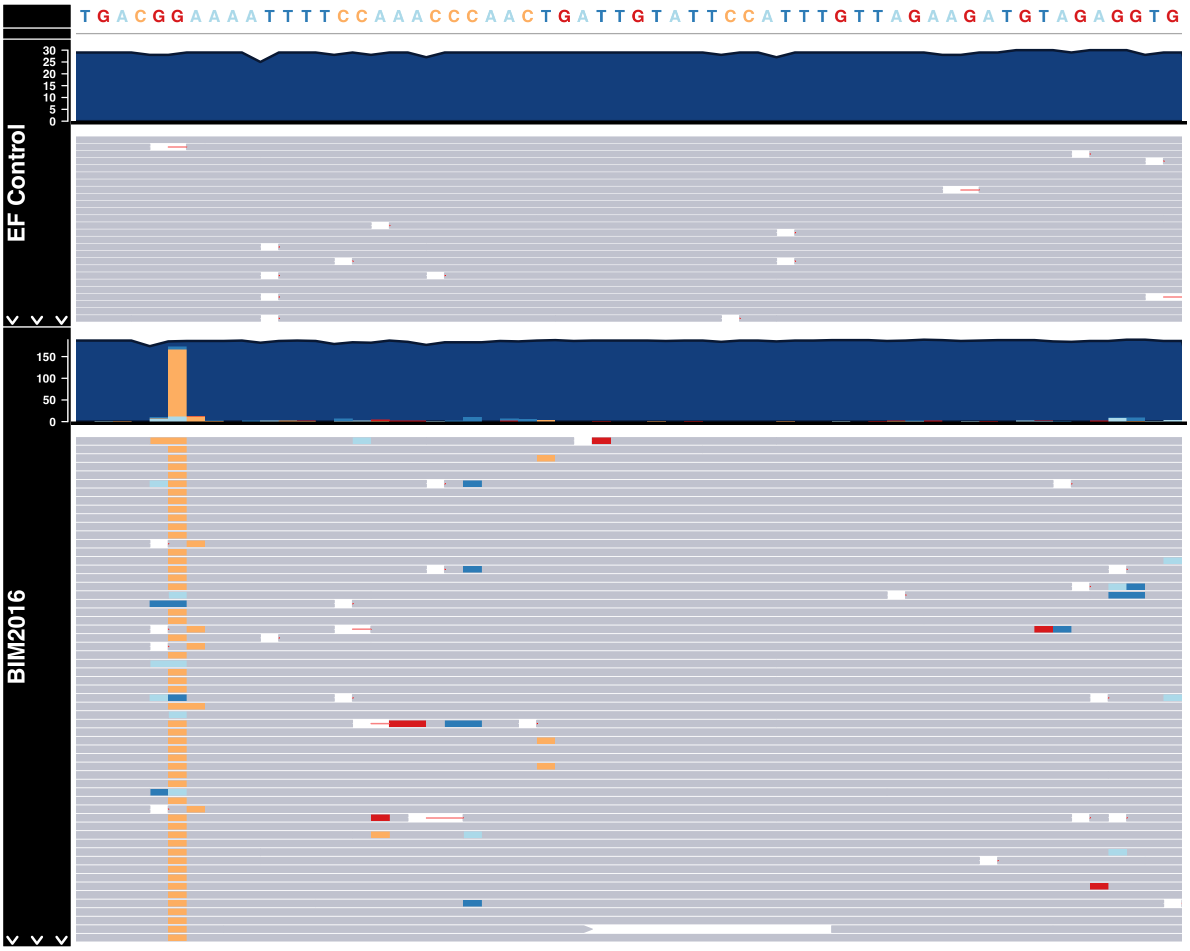


**Figure S 12.** Pileup plot showing the T->A SNV in wcaG gene in BIM-2016 compared to the reference genome ATCC-700802. In each panel, the read coverage is shown in blue, each mapped read is represented in grey, with light blue indicating the variant. The control (ATCC-700802) is shown in the top panel, while BIM20 is shown below.

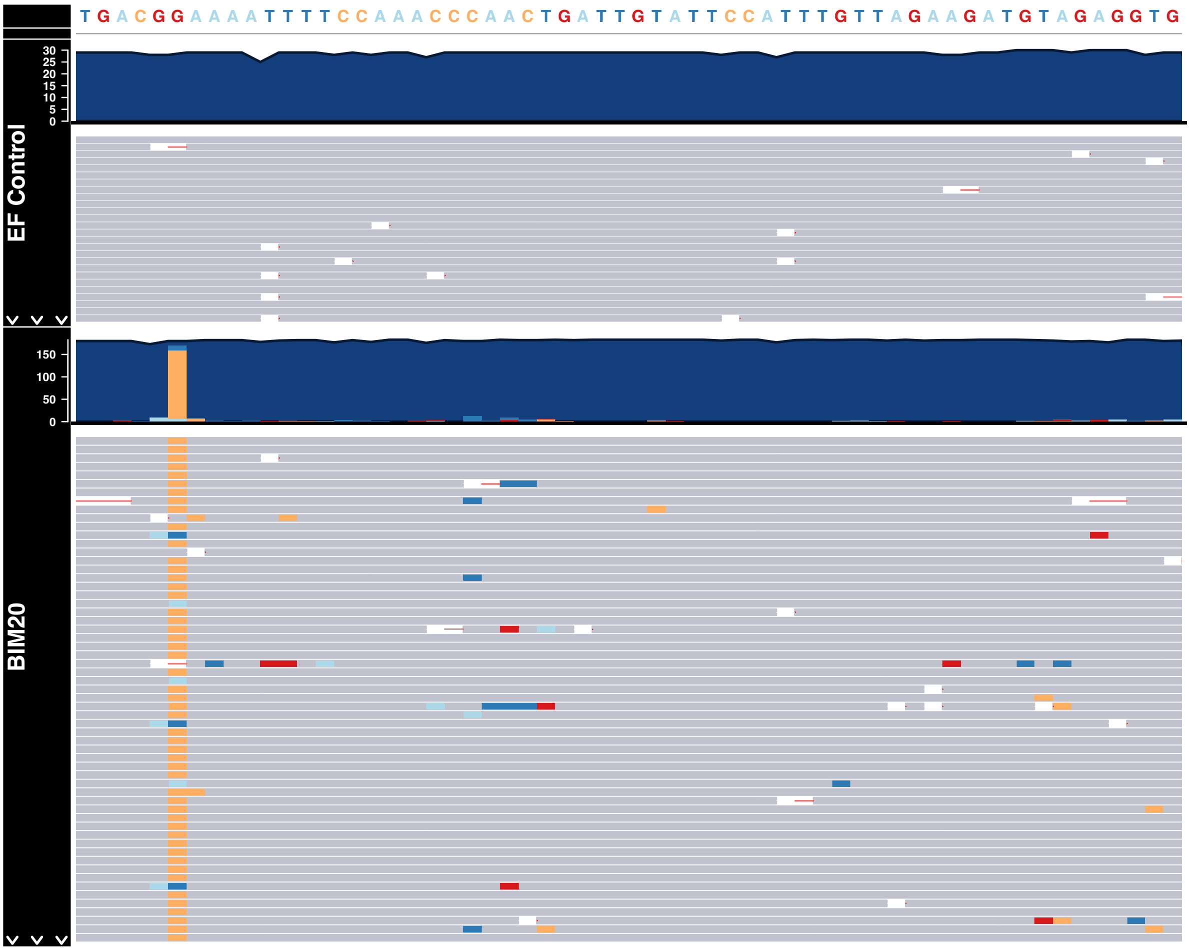


**Figure S 13.** Pileup plot showing the G->C SNV 109 bases upstream of LicD in BIM-20 compared to the reference genome ATCC-700802. In each panel, the read coverage is shown in blue, each mapped read is represented in grey, with yellow indicating the G->C variant. The control (ATCC-700802) is shown in the top panel, while BIM-20 is shown below.

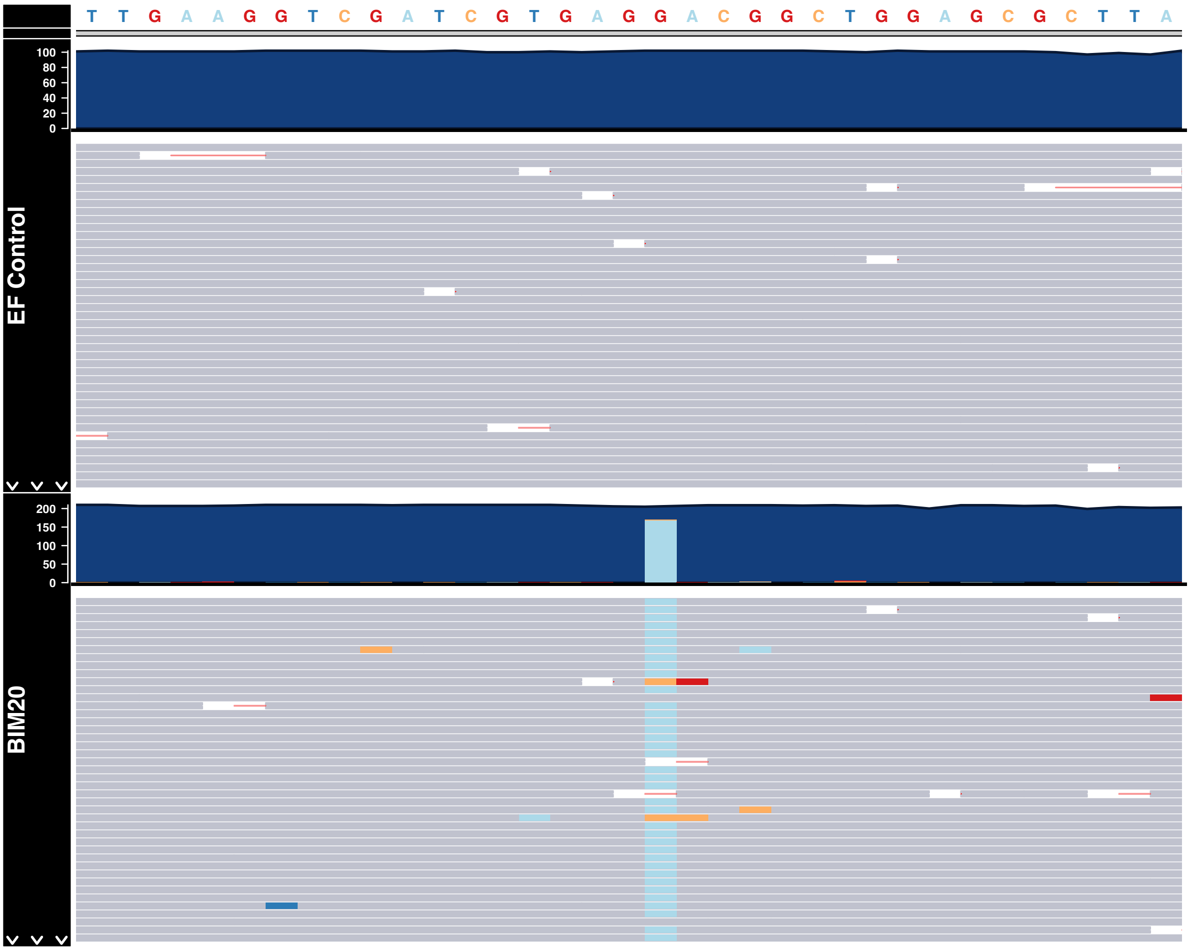


**Figure S 14.** Pileup plot showing the C->A SNV in 23S rRNA gene in BIM-20 compared to the reference genome ATCC-700802. In each panel, the read coverage is shown in blue, each mapped read is represented in grey, with light blue indicating the variant. The control (ATCC-700802) is shown in the top panel, while BIM-20 is shown below.

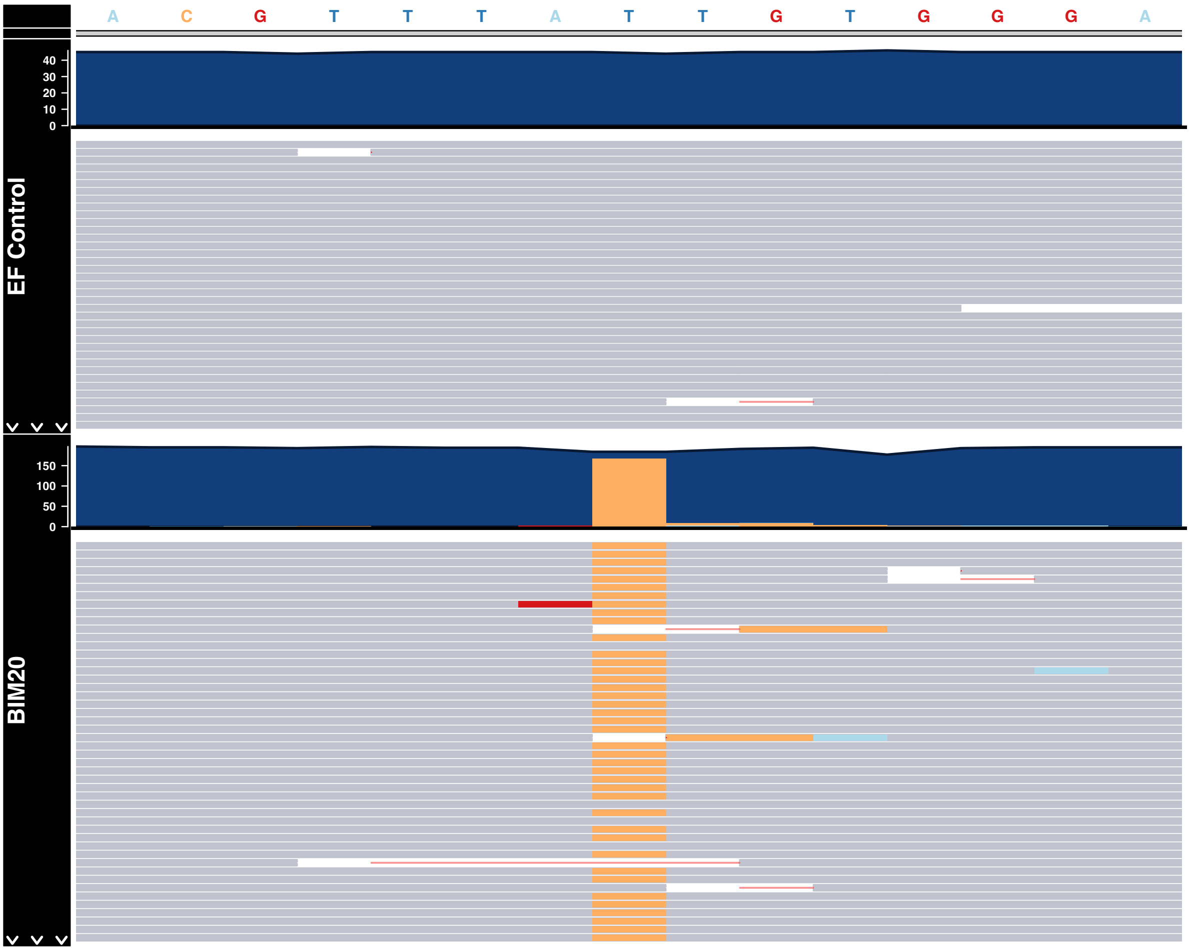


**Figure S 15.** Pileup plot showing the T->C SNV in FabZ gene in BIM-20 compared to the reference genome ATCC-700802. In each panel, the read coverage is shown in blue, each mapped read is represented in grey, with yellow indicating the variant. The control (ATCC-700802) is shown in the top panel, while BIM-20 is shown below.

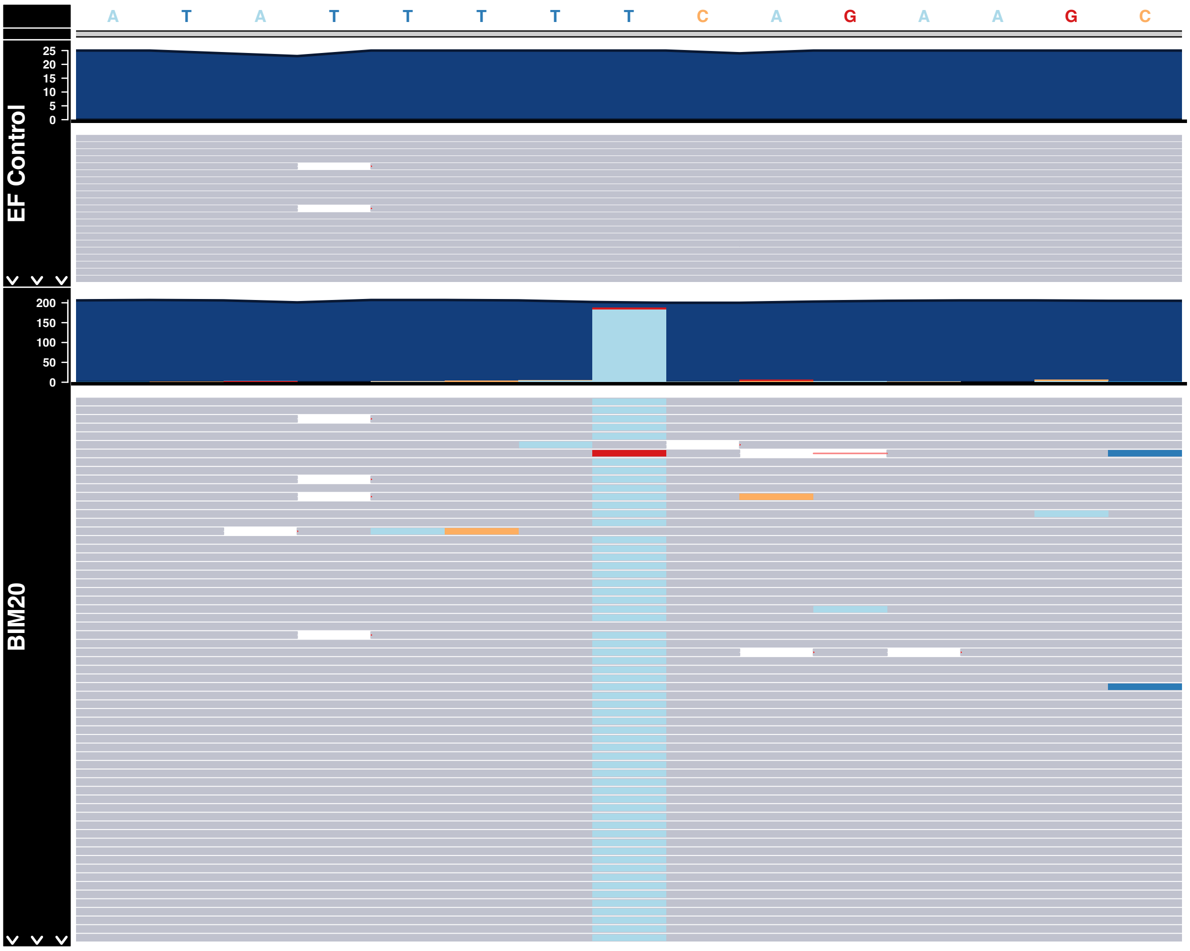


**Figure S 16.** Pileup plot showing the T->A SNV in wcaG gene in BIM-20 compared to the reference genome ATCC-700802. In each panel, the read coverage is shown in blue, each mapped read is represented in grey, with light blue indicating the variant. The control (ATCC-700802) is shown in the top panel, while BIM-20 is shown below.

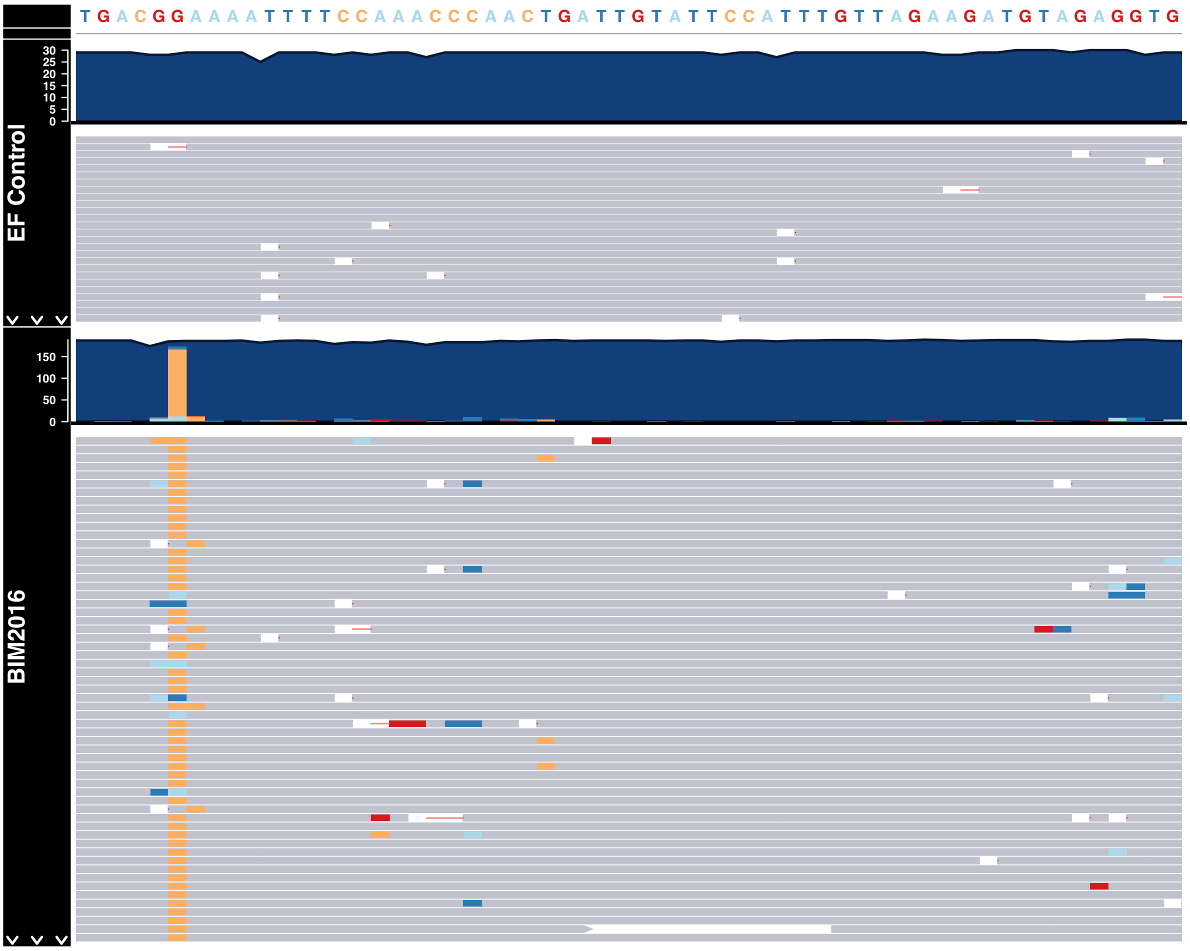


**Figure S 17.** Pileup plot showing the G->C SNV 109 bases upstream of LicD in BIM-2016 compared to the reference genome ATCC-700802. In each panel, the read coverage is shown in blue, each mapped read is represented in grey, with yellow indicating the G->C variant. The control (ATCC-700802) is shown in the top panel, while BIM-2016 is shown below.

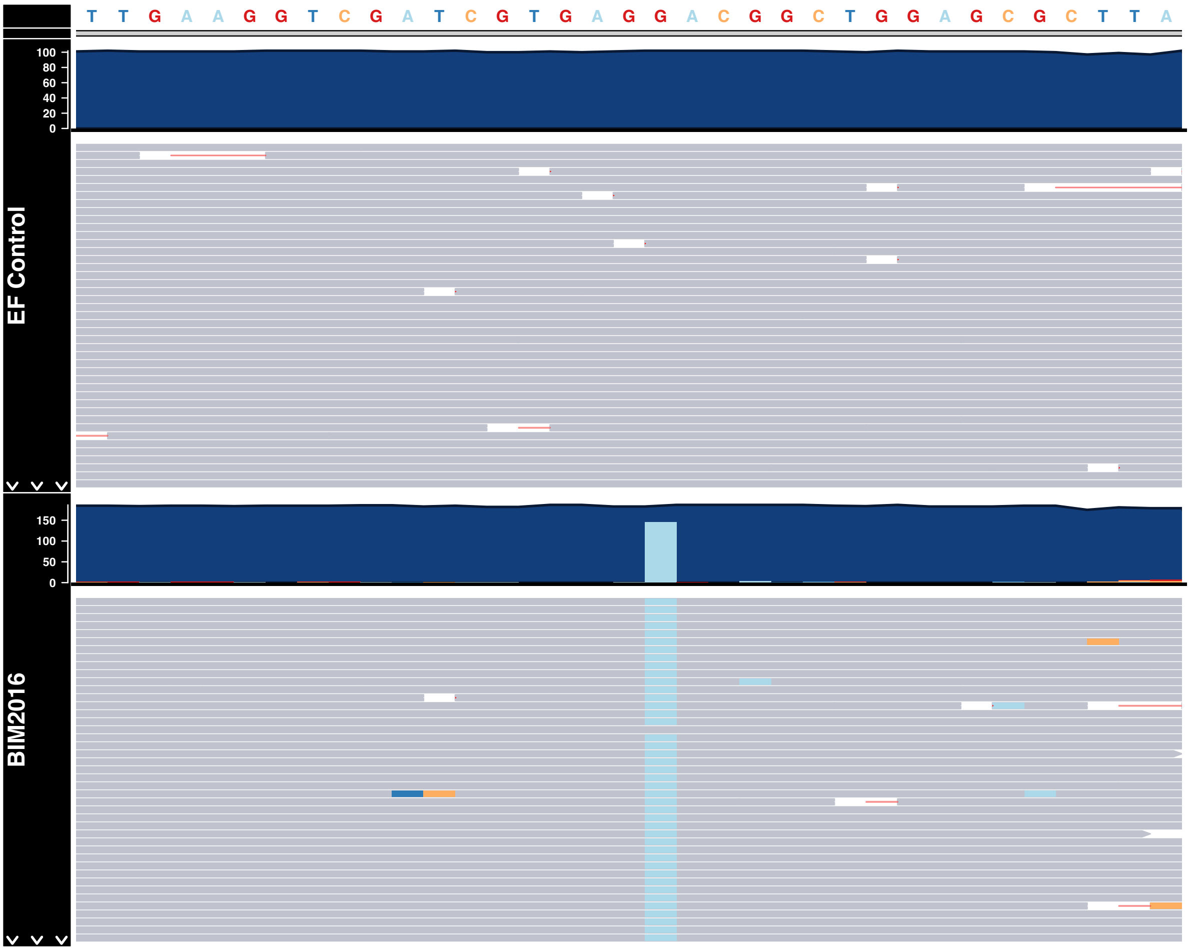


**Figure S 18.** Pileup plot showing the C->A SNV in 23S rRNA gene in BIM-2016 compared to the reference genome ATCC-700802. In each panel, the read coverage is shown in blue, each mapped read is represented in grey, with light blue indicating the variant. The control (ATCC-700802) is shown in the top panel, while BIM-2016 is shown below.

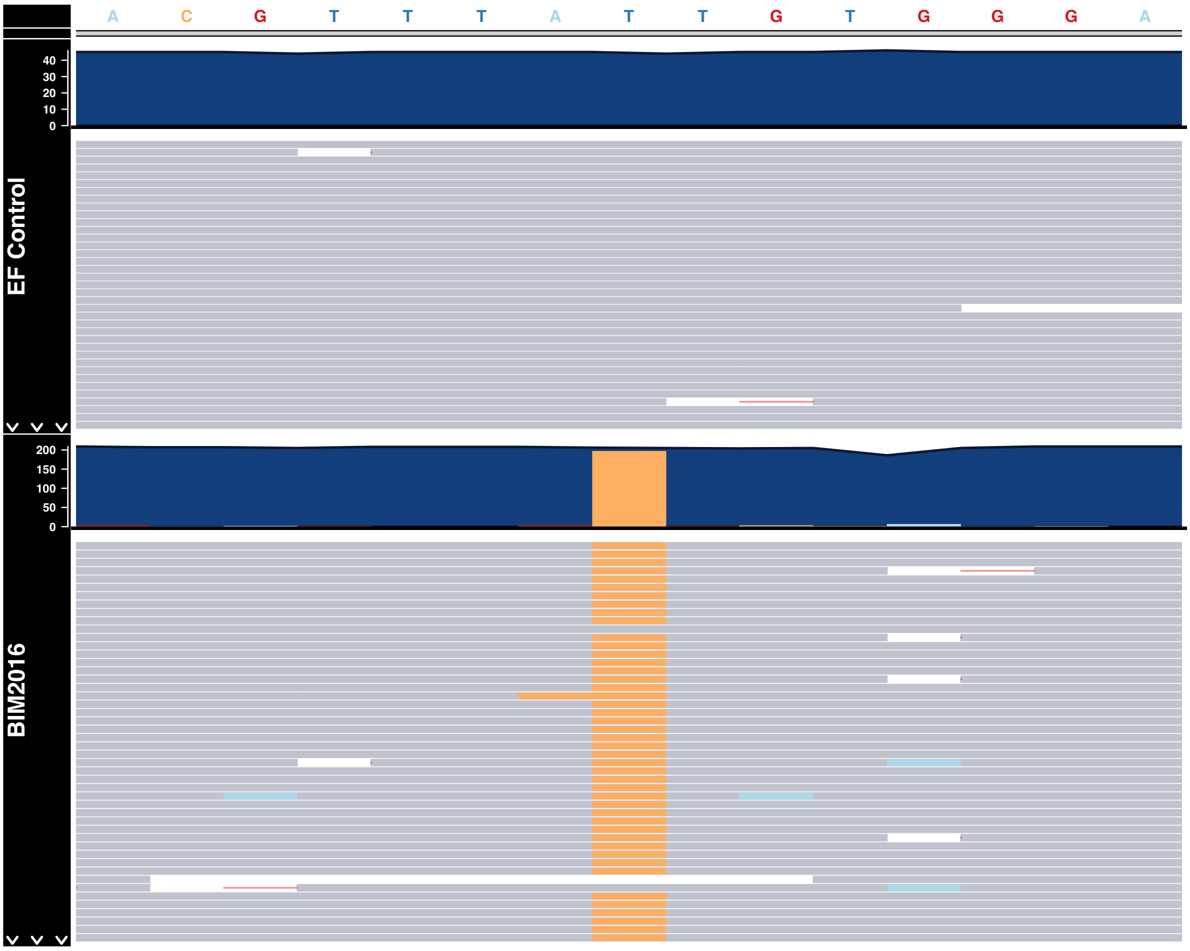


**Figure S 19.** Pileup plot showing the T->C SNV in FabZ gene in BIM-2016 compared to the reference genome ATCC-700802. In each panel, the read coverage is shown in blue, each mapped read is represented in grey, with yellow indicating the variant. The control (ATCC-700802) is shown in the top panel, while BIM-2016 is shown below.

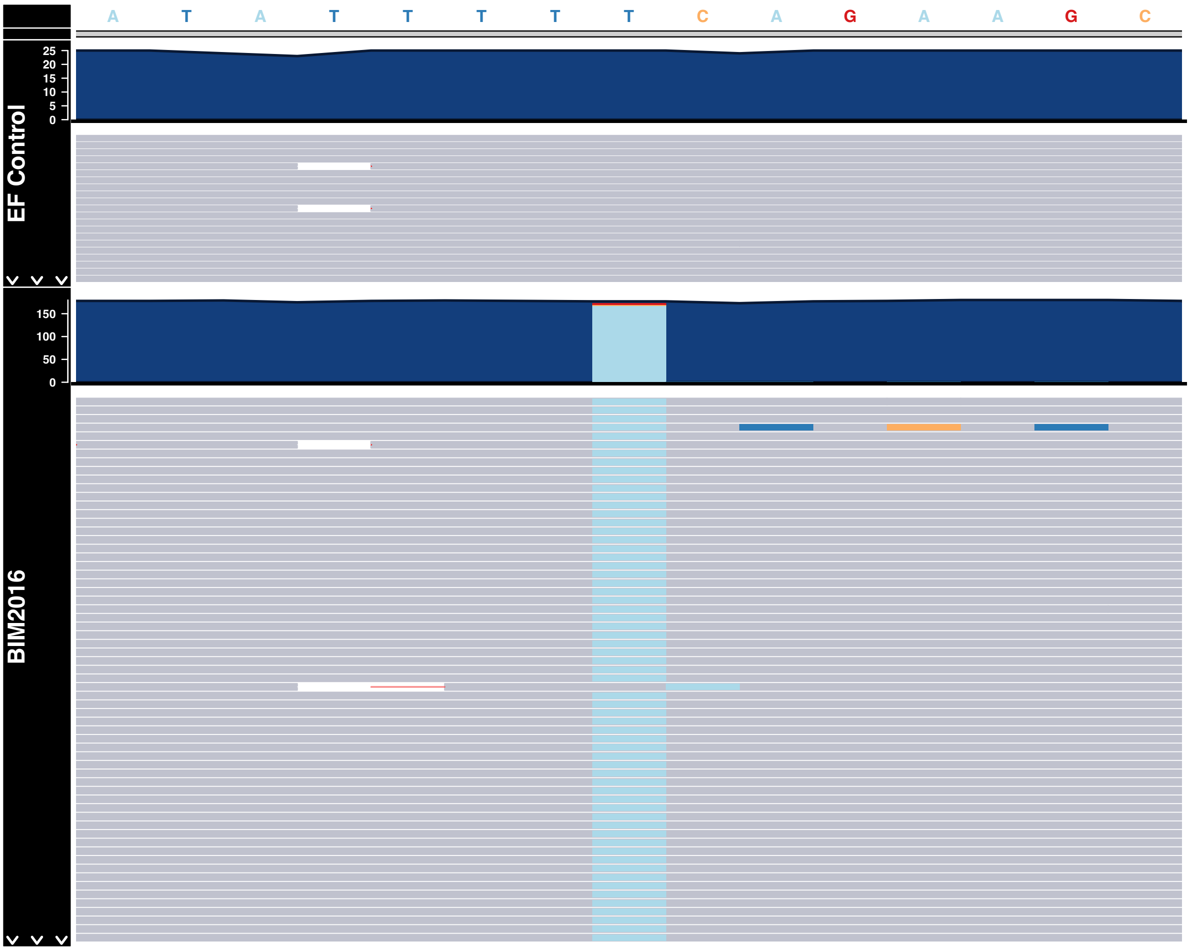


**Figure S 20.** Pileup plot showing the T->A SNV in wcaG gene in BIM-2016 compared to the reference genome ATCC-700802. In each panel, the read coverage is shown in blue, each mapped read is represented in grey, with light blue indicating the variant. The control (ATCC-700802) is shown in the top panel, while BIM-2016 is shown below.
